# Of Biofilms and Beehives: An Analogy-Based Instructional Tool to Introduce Biofilms to High-School and Undergraduate Students

**DOI:** 10.1101/2021.10.27.466040

**Authors:** Snehal Kadam, Ankita Chattopadhyay, Karishma S Kaushik

## Abstract

The concept of biofilms and biofilm-based research is largely absent or minimally described in high school and undergraduate life science curriculum. While it is well-established that microbes, such as bacteria and fungi most often exist in multicellular biofilm communities, descriptions in standard biology textbooks continue to focus on the single-celled form of microbial life. We have developed an analogy-based instructional tool to introduce and explain biofilms to high school and undergraduate students. The module employs an analogy with beehives, given that biofilms and beehives are both ‘superorganism’ states, to explain key biofilm features such as development and structure, chemical communication, division of labor and emergent properties. We delivered this analogy based learning tool to a cohort of 49 high school and undergraduate students, and based on participant feedback and learnings, present a formal evaluation of the instructional tool. Further, we outline prerequisites and learning approaches that can enable the delivery of this module in classroom and virtual learning settings, including suggestions for pre-lesson reading, student-centred interactive activities, and specific learning objectives. Taken together, this instructional analogy holds potential to serve as an educational tool to introduce biofilms in high school and undergraduate curricula in a relatable and comprehensible manner.

## INTRODUCTION

Typically studied as single-celled organisms, microbes are known to form large multicellular communities known as biofilms (1–3). Biofilms are highly-organized, structured, three-dimensional, aggregates of microbial cells (bacteria or fungi), embedded in a self-produced extracellular matrix (4). Biofilms have serious health consequences, given their implication in a range of human infections (5–9), as well as affect the environment via water pollution and industrial fouling (10, 11). While biofilms are actively-studied in research laboratories, they largely absent or minimally described in school and undergraduate biology curricula (12). For example, the high school textbook Life on Earth (13) discusses microbial communities in the context of biodiversity, but does not discuss the biofilm mode of microbial life. On the other hand, the widely-used undergraduate biology textbook Concepts of Biology (14) includes a very brief overview of biofilms, with no insights into the processes involved in biofilm structure, formation and function. Given this, an instructional tool that introduces and discusses biofilms in a relatable and comprehensible manner could serve as a valuable addition to high school and undergraduate biology curricula.

Analogies have been explored as tools in science education (15–19), and the use of analogies is based on deconstructing systems with respect to their parts, the relation between the parts and the agreement across the themes (16, 19–26). Given this, analogical teaching is inherently suited for building new concepts, and holds value in introducing new concepts in an engaging and relatable manner (19, 27, 28). To develop an analogy-based instructional tool for biofilms, we use an analogy with beehives, given that biofilms and beehives are both collective organismal states, also known as ‘superorganisms’ (29, 30). Beehives are macroscopic, visible to the naked eye, and well-known entities, and provide a relatively familiar analog for microscopic, multicellular microbial biofilms. Using an analogy with beehives, this instructional tool introduces and explains key biofilm features such as development and structure, chemical communication, division of labor, and emergent properties. Since a large segment of research on biofilms has focused on bacteria, we have largely drawn examples from bacterial biofilms.

### Intended audience

This analogy based instructional tool is intended for high school and undergraduate students, as part of the biology or life science curriculum. The content draws from fields such as biology, microbiology, chemistry and biochemistry, and can be adapted (as per suggestions provided) for the different levels and.

### Learning time

The entire analogy-based lesson, including delivery of the content, pre-and post-session feedback and student activities, requires 90 minutes. The instructor leads the delivery of the content, which is facilitated by student learning activities (as individuals and in groups). The curriculum tool includes instructor guidelines for delivery and student activities, slides for delivery, and sample feedback forms.

### Prerequisite student knowledge

High school students and undergraduates would be expected to have had a science or biology course. No prior laboratory of field experience is required. To bring students on the same level, the delivery of the analogy includes a set of pre-session reading materials, namely relevant chapters in biology text books (recommended chapters include basics of biology, biological communities, chemistry of life, cellular life, DNA and genes, diversity of life and bacterial life, as well as a list of 24 key words, across the concepts of biofilms and beehives (Suppl Material). These can be provided to the students a week prior to the session.

### Learning objectives

The learning objectives for this instructional tool are as follows:

1. At the end of the lesson, students will be able to recognize the concept of biofilms as bacterial communities, and contrast it from single-celled microbial life.
2. On completion of the lesson, students will be able to identify the importance of studying biofilms, from both the health and environmental perspective.
3. From the section on development and structure, students will be able to identify and recognize the typical structure of biofilms, the five main stages in biofilm formation, and events influencing these stages.
4. After reading the section on chemical communication, students will be able to recognize the phenomenon of quorum sensing, and the roles of different autoinducer molecules.
5. On completion of the section on division of labor, students will be able to recapitulate the example of division of labor in *B. subtilis* biofilms that contributes to the formation of the biofilm extracellular matrix.
6. Based on their understanding of emergent properties, students will be able to identify the definition of emergent properties, and recognize why antibiotic tolerance is a feature of multicellular biofilms (as opposed to single cells).
7. After the section on limitations of the analogy, students will be able to contrast the key areas in which biofilms are different from beehives by listing at least one key difference.
8. Students will be able to apply their understanding of the analogy to develop at least one idea of their own related to a new idea of investigation on biofilms. It is important to note that here ‘new’ represents what is not stated or explained in the analogy, and not new for the field per se, given that students may not have a comprehensive overview of the current status of biofilm research.

## PROCEDURE

### Materials

Delivery of the analogy-based instructional tool will require a classroom setting (if in-person) or a virtual platform. If in person, the instructor will require equipment for projection of slides (overhead projector, computer). Students will require writing tools and sheets of paper for notes.

### Student instructions

Prior to the lesson, students would be expected to read the suggested pre-session reading materials such as select textbook chapters, as well as familiarize themselves with the meanings of the key words provided to them.

### Faculty instructions

Prior to the lesson, the faculty instructor will need to share the suggested pre-session reading materials and key words with the students. The instructor will also need to read and understand the modules of the analogy (Table 1), and download and familiarise themselves with the slide deck (Suppl Material) provided for delivery of the analogy. The module includes references to original scientific literature which may be used by the instructor to further clarify concepts. A detailed guideline for the delivery of the analogy, including time to be allocated for each section of the module and additional learning activities, is provided in the Supplementary Material.

**Table 1:** Summary of key shared and non-shared features related to the analogy on biofilms and beehives.

| Features | Beehives | Biofilms |
| --- | --- | --- |
| **Development and Structure** | Self-produced beeswax<br>Structured solid (at room temperature) beeswax<br>Defined 3D hexagonal structure<br>Honeycomb has mechanical and protective role<br>‘Scouting behaviour’ followed by mass recruitment | Self-produced EPS matrix<br>Slimy, hydrated, dynamic matrix<br>Variable structure, shape, and 3D architecture<br>EPS matrix has mechanical, protective and transport roles<br>Initial reversible attachment followed by irreversible phase |
| **Chemical Communication** | Self-produced pheromones<br>Different types of pheromones based on bee species and function<br>Controls behaviours such as colony development, reproduction, brood rearing<br>Mainly for intraspecies communication, rare pheromones for interspecies communication | Self-produced autoinducers<br>Different types of autoinducers based on microbial species<br>Controls behaviours such as biofilm formation, virulence, bioluminescence<br>Autoinducers for Intraspecies, interspecies and interkingdom communication |
| **Division of Labor** | Presence of different subpopulations of a single species<br>Subpopulations have committed lifestyle and function | Presence of different subpopulations consisting of single and mixed species<br>Role and function subpopulations may vary depending on genetic and environmental changes |
| **Emergent Properties** | Emergent properties result from an active response (shivering in bees leading to thermoregulation of the hive) | Emergent properties typically result from the presence of passive structures (physical matrix barrier leading to antibiotic tolerance) |

### Module 1: Development and structure of beehives and biofilms

The Western or European honeybee (*Apis mellifera*) lives in well-ordered colonies or beehives, consisting of a large queen bee, thousands of female worker bees, which lack a completely developed reproductive system, and a few drone males (31–33). Worker bees make the 3D honeycomb structures of hives from beeswax secreted from their abdominal glands (34). At the start of building the colony, scout bees search for suitable locations to occupy, known as ‘scouting behaviour’, based on criteria such as position and volume of the nest cavity, light, moisture, and temperature (35–37). Scout bees communicate the presence of an optimum location via a characteristic ‘waggle dance’ (38), and if several bees come back with the same information, the waggle dance spreads to other bees. When one site is being visited by a sufficiently large number of scouts (39), the recruitment dance results in a swarm, where the queen bee and scouts proceed to establish a colony at the chosen site (40) (Figure 1A).

**Figure 1:**
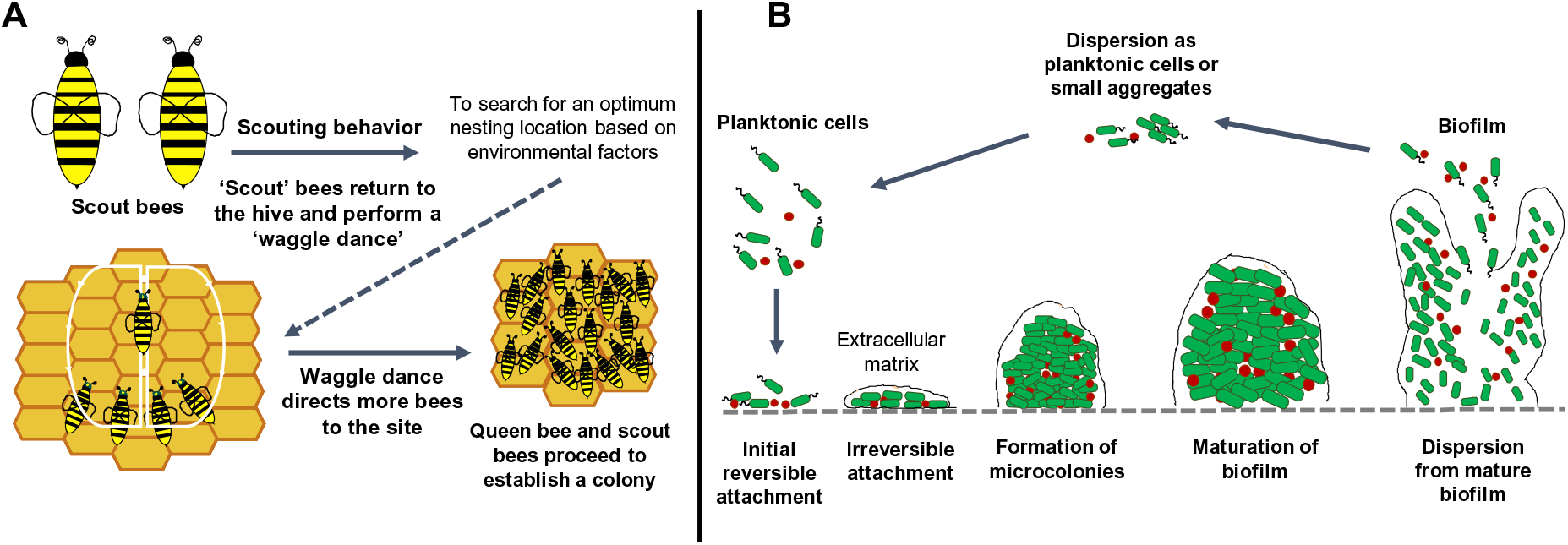
Development and structure of beehives and biofilms, illustrating shared and non-shared features between the two entities. In *Apis mellifera*, a few scout bees search for an optimum nesting location, the presence and location of which is communicated to the hive using a ‘waggle dance’, among other characteristic movements. These movements direct more bees, including the Queen bee, to the site, which proceed to establish a new colony. In bacterial biofilms, a few planktonic cells or aggregates proceed to initiate biofilm formation following stages of initial reversible and then irreversible attachment, to biotic or abiotic surfaces. Following this, attached bacteria grow to form microcolonies and three-dimensional structures, constituting a mature biofilm. Bacterial cells disperse, as small aggregates or single cells, and can proceed to establish biofilm formation at other sites. Illustrated here is a typical mixed-species biofilm with two different bacterial species shown in green (bacilli) and red (cocci).

In the early stages of biofilm formation, free-floating bacteria (or small bacterial aggregates) attach to a biotic or abiotic surface, via weak chemical forces (41). This initial attachment of free-floating bacteria is influenced by factors such as temperature, surface properties and nutrients (41). At this early stage, bacteria interact with the surface in a transient manner, with each reversible contact priming the bacteria for the next stage of irreversible attachment (41). In the presence of high bacterial densities, these transiently attached bacterial cells secrete extracellular polymeric substance (EPS) (42, 43). EPS consists of polysaccharides, proteins, and extracellular DNA, and helps bacteria adhere to surfaces and provides mechanical strength to biofilms (41, 44). During further maturation, the biofilm extends from the surface to develop multiple layers of bacterial microcolonies, thereby building a 3D structure. The EPS matrix is a dynamic substrate that influences the transfer of nutrients and metabolites, resulting in chemical gradients in the biofilm structure (45–47). Finally, a mature biofilm can disperse, either as clumps of cells or single cells, to seed new surfaces (Figure 1B).

### Module 2: Chemical communication in beehives and biofilms

Chemical communication in beehives occurs through pheromones, secreted from exocrine glands (48). Honeybee pheromones are a mixture of volatile and non-volatile chemical substances that are transmitted by direct contact. Specific pheromones are released by queen, worker, drone and brood bees, ensuring a broad range of functions (48). The queen signal is a complex mixture of several chemicals, the main component being Queen Mandibular Pheromone (QMP). QMP is responsible for worker activities, drone attraction and queen rearing. Worker bees produce Nasonov gland pheromones to drive the returning forager bees back to the hive, mark hive entrance, locate food resources and rear future queens (49). Other important pheromones include alarm pheromones, drone pheromones, which are almost exclusively linked to mating functions, and brood pheromones, that regulate colony development and formation (48) (Figure 2A).

**Figure 2:**
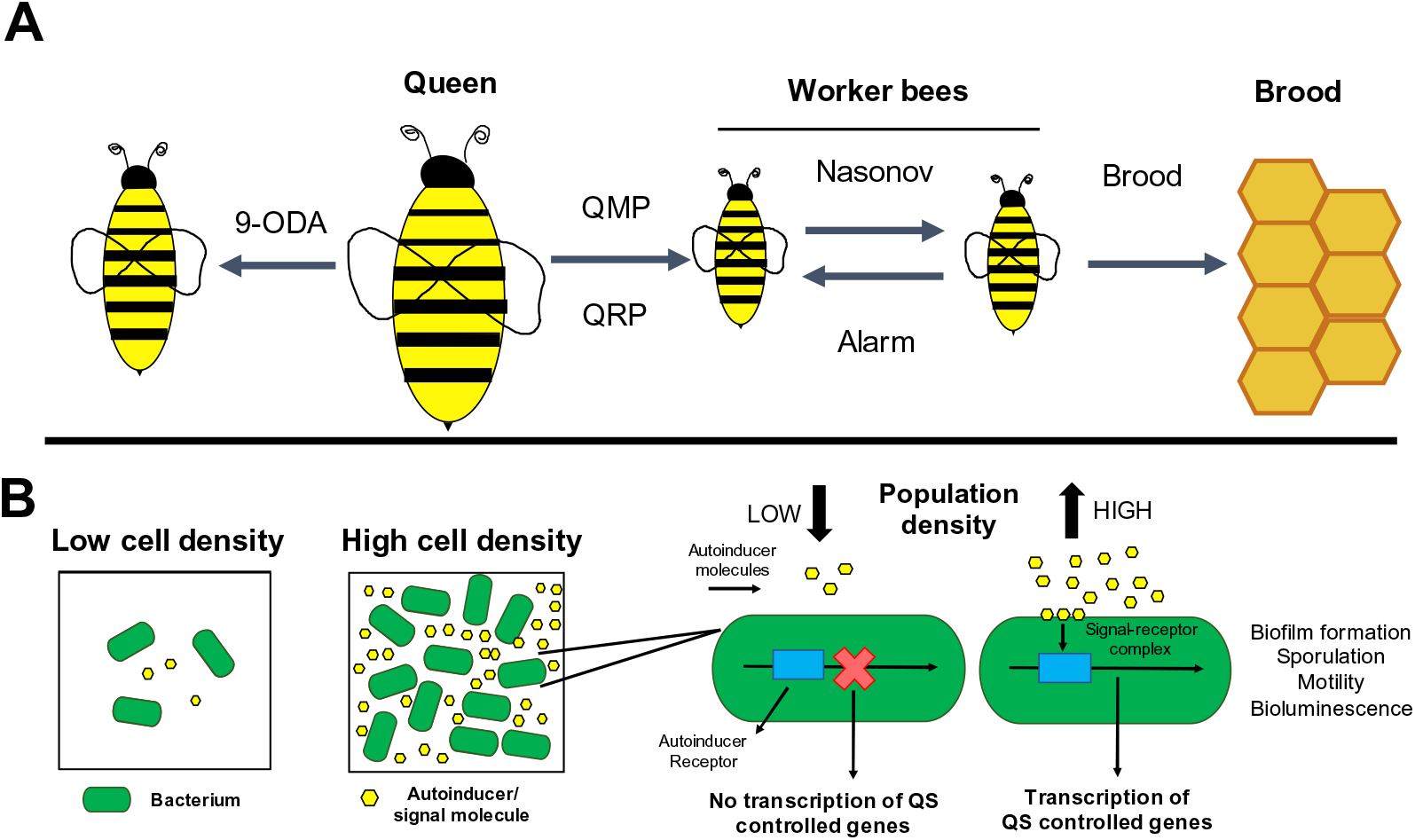
Chemical communication in beehives and biofilms, illustrating shared and non-shared features between the two entities. In honeybees (such as *Apis mellifera*), chemical communication occurs through specific pheromones, volatile and non-volatile chemical substances transmitted by direct contact. Released by queen, worker, drone and brood bees, pheromones are responsible for a range of functions such as colony development and formation, location of food resources, drone attraction and queen and brood rearing. Similar to the chemical communication in bees, bacteria communicate via signalling molecules (such as autoinducers), in a density-dependent process. At high bacterial cell densities, the signal concentration reaches a certain threshold, following which autoinducer molecules bind to bacterial cell surface receptors, triggering population-wide changes in gene expression. These changes regulate various bacterial group functions such as biofilm formation, motility, sporulation, and virulence.

Similar to the chemical communication in bees, bacteria communicate via signalling molecules (50–52). This phenomenon, known as quorum sensing, depends on bacterial cell density and is mediated by small, diffusible extracellular signal molecules or autoinducers (53). As the local density of the bacteria increases, the extracellular concentration of autoinducers also increases (Figure 2B). Autoinducer molecules bind to bacterial cell receptors, triggering changes in gene expression across the bacterial population (54). These changes in gene expression regulate the various stages of biofilm formation (50, 51, 55, 56). There are three major classes of autoinducer molecules, N-acylated homoserine lactones (AHLs) or autoinducer-1 (AI-1) primarily found in Gram-negative bacteria, oligopeptides found in Gram-positive bacteria, and autoinducer-2 (AI-2), which is present in Gram-positive and Gram-negative bacteria, and enables communication across bacterial species (54, 57, 58).

### Module 3: Division of labor within beehives and biofilms

Honeybees exhibit a unique haplodiploid mode of sex-determination in which the unfertilized haploid eggs produce males, while worker and queen females hatch from the fertilized diploid eggs (59–61). Different subpopulations of bees exist within the hive, with each subpopulation performing specialized functions to maintain the hive integrity (Figure 3A). Typically, there is only one queen bee per colony, who is fertile and lays eggs in the hive. Drone bees are the sole males of the colony, and their main task is to fertilize a receptive queen. Worker bees do the majority of the work for the colony, and there is further division of labor within them (29, 31, 62). Activities divided among worker bees include nursing the developing larvae and the queen, cleaning and building the hive, foraging for pollen, storing honey and nectar in the hive, and protecting the hive from predators.

**Figure 3:**
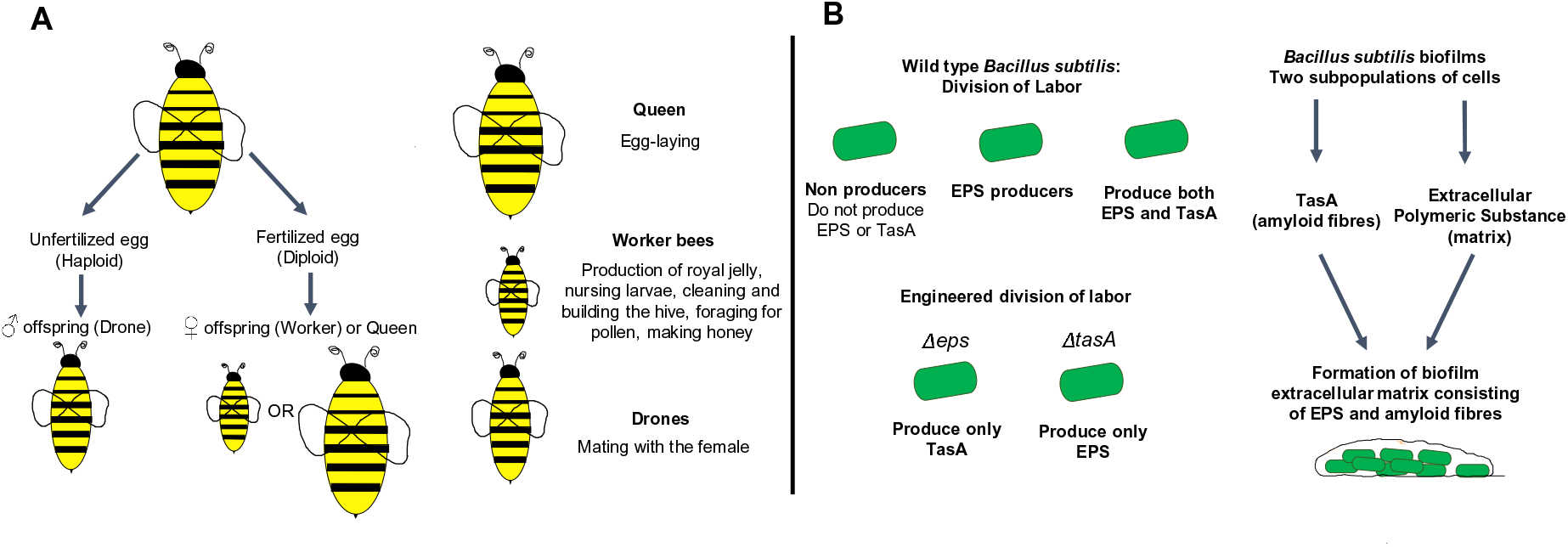
Division of labor in beehives and biofilms, illustrating shared and non-shared features between the two entities. Honeybees exhibit a unique haplodiploid mode of sex-determination in which the unfertilized haploid eggs produce males, while worker and queen females hatch from the fertilized diploid eggs. In honeybees (such as *Apis mellifera*), these different subpopulations of bees perform specialised functions. Typically, there is only one queen bee per colony, who is fertile and lays eggs in the hive. Drone bees are the sole males of the colony, and their main task is to fertilize a receptive queen. Worker bees do the majority of the work for the colony, with further division of labor within them. A form of division of labor, based on differential gene expression in the bacterial population, is also observed in biofilms formed by the bacterium *Bacillus subtilis*, where division of tasks between subpopulations contributes to the formation of the biofilm matrix.

A form of division of labor, based on differences in gene expression in the population, is also observed in bacterial biofilms (63–67). For example, in *Bacillus subtilis* biofilms, there is division of tasks between groups of populations that contributes to the formation of the biofilm matrix (65). There are two main constituents of the matrix namely an extracellular polymeric substance (EPS) and a protein TasA (amyloid fibres), which are produced by different *B. subtilis* subgroups. Cells within the biofilm segregate into groups that either produce both components, produce EPS only or produce neither (Figure 3B). When mutants *Δeps* (producing only the TasA protein) and *ΔtasA* (producing only EPS) were mixed in a culture, they complemented each other by sharing EPS and TasA, to make a biofilm similar to the wild-type (with no mutations). However, these mutants, when studied individually, were deficient for biofilm formation.

### Module 4: Emergent properties in beehives and biofilms

As ‘superorganism’ states, beehives and biofilms exhibit collective or emergent properties, that are not displayed by individual organisms (68). In honeybee colonies, a well-known emergent behaviour is thermoregulation (69), that relates to the ability of the honeybee colony to survive as a whole. At low temperatures, bees tend to move closer together and share body heat. Since the centre has more heat, and younger bees cannot shiver, they move inwards. Adult bees shiver to produce heat and move to the middle and outer layers. This heat warms the whole hive (Figure 4A). As the heat in the centre increases leading to a situation of excess heat, the young bees move to create channels of air exchange, allowing heat from inner regions to flow out towards the older bees. This combined effect enables the hive as a whole to stay warm, a critical factor for survival of the colony.

**Figure 4:**
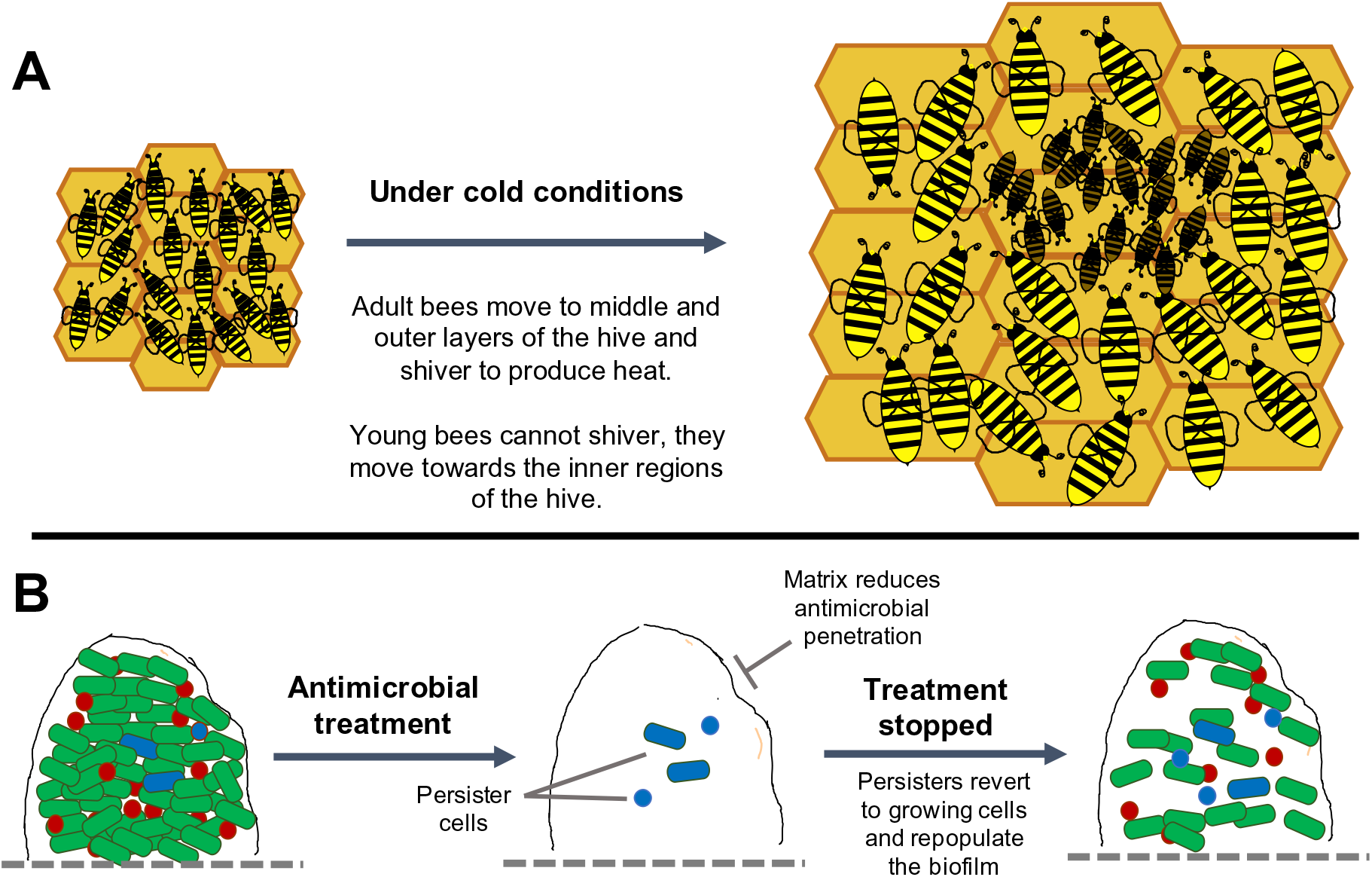
Emergent features in beehives and biofilms, illustrating shared and non-shared features between the two entities. In honeybee colonies, a well-known emergent behaviour is thermoregulation, which relates to the ability of the honeybee colony to survive as a whole. At low temperatures, bees tend to move closer together and share body heat. Since the centre has more heat, and younger bees cannot shiver, they move inwards. Adult bees shiver to produce heat and move to the middle and outer layers. In biofilms, an important emergent property is the increased tolerance to antimicrobial treatments. This results from various factors such as physical and chemical factors in the EPS that reduce the diffusion of antimicrobial agents into the inner parts of the biofilm, as well as the metabolic heterogeneity of the biofilm state that contributes to the formation of slow-growing or dormant cells, that can tolerate high concentrations of antibiotics, and can repopulate the biofilm once antimicrobial treatments are stopped.

An important emergent property of bacterial biofilms, that differs from free-floating cells, is increased tolerance to antimicrobials. This results from specific properties of bacteria in the biofilm, as well as the biofilm matrix itself (2). The EPS matrix reduces diffusion of antimicrobial agents into the inner parts of the biofilm (2). In the bacterium *Pseudomonas aeruginosa,* components of EPS such as polysaccharide and extracellular DNA, form interactions with antibiotics and impede their penetration through the matrix (70). Bacteria in biofilms also adopt properties of slow growth and dormancy, with reduce their susceptibility to antibiotics, such as penicillin, that act on actively-growing cells (71). One type of slow-growing cells in biofilms are persister cells, that exhibit high-level tolerance to antimicrobials (72, 73), but can revert to a growing state and repopulate the biofilm once treatment is stopped (Figure 4B).

### Suggestions for determining student learning

Student learning was assessed using pre-session and post-session feedback forms via Google forms. The forms used a combination of multiple choice and free response questions. Pre-session feedback included information on participant demographics, prior science and biology courses, use of pre-session reading materials and familiarity with brood concepts of the analogy. Post-session feedback assessed learning of the content delivered in the modules and stated learning objectives. Students were provided time to fill these forms before and after the session. Both feedback forms are available in the Supplementary Material.

### Sample Data

In the pre-session form, students provided data related to demographics, educational level, previous science or biology courses, familiarity with biofilms and beehives and use of pre-session reading materials (Figures 5-7). Students learning data was collected in response to specific content-based questions, anecdotal feedback in response to open-ended questions, and new ideas and hypotheses generated from the modules (Figures 8-11 and Tables 2-5).

**Figure 5:**
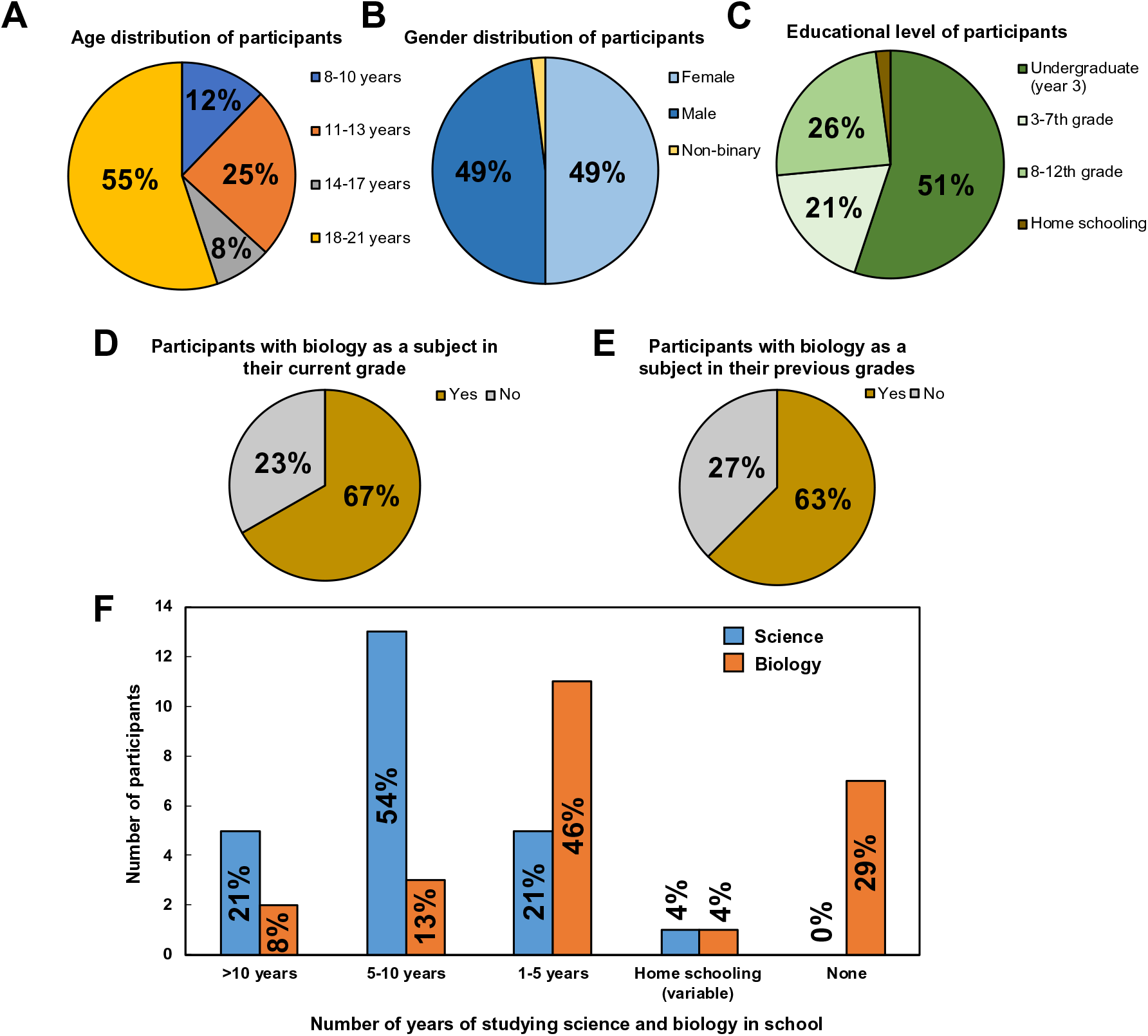
Distribution of participants based on age, gender and educational level. The analogy-based session was delivered to a total of 49 students (24 high school students and 25 undergraduates) across India. Participants were pursuing schooling across different systems (state, central home schooling), and reported varied years of science and biology education (*n=49 for 5A, B, C and n=24, for 5D, E, F)*.

**Figure 6:**
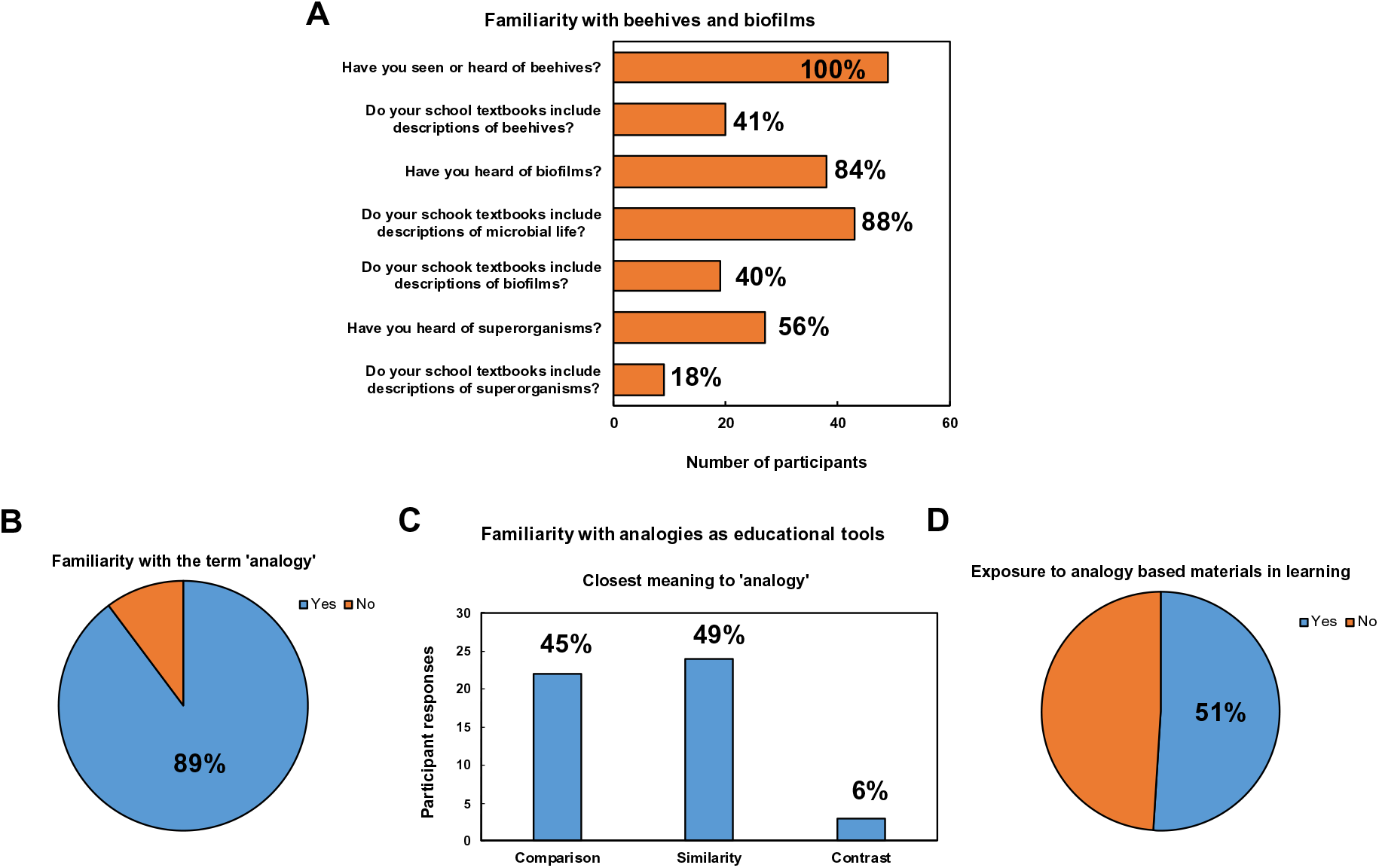
Prior familiarity of participants with the key concepts in this analogy-based lesson. Participants reported familiarity with key concepts such as beehives, biofilms and superorganisms, however, biofilms were notably less represented in school and undergraduate textbooks (*n=49)*.

**Figure 7:**
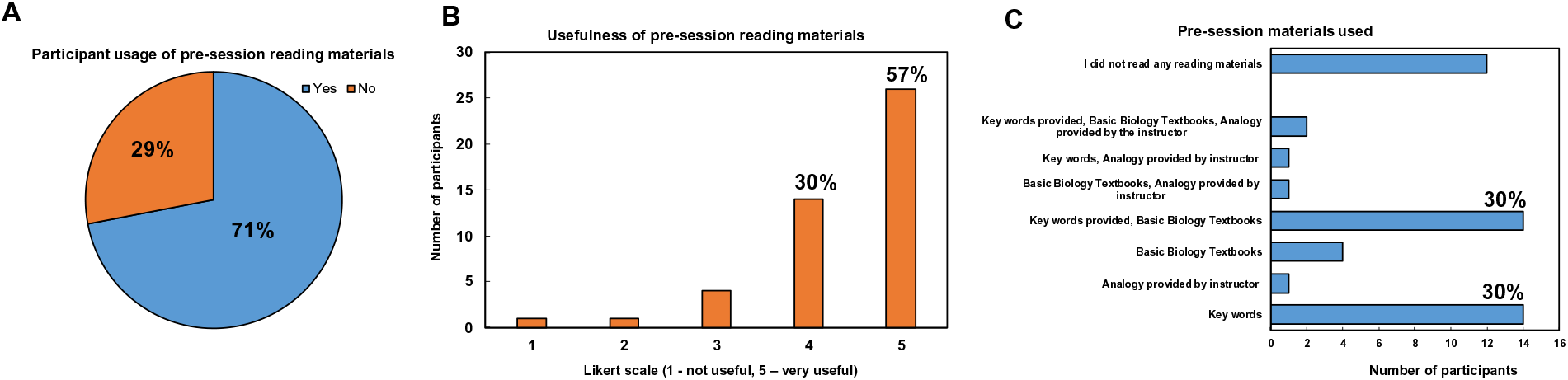
Participant usage of pre-session reading materials. Prior to the delivery of analogy-based session, participants were provided a set of pre-session reading materials which included key words, relevant concepts in basic biology textbooks and the draft of the analogy. Based on feedback, these materials were both utilized and reported to be useful by the respondents (*n=49)*.

**Figure 8:**
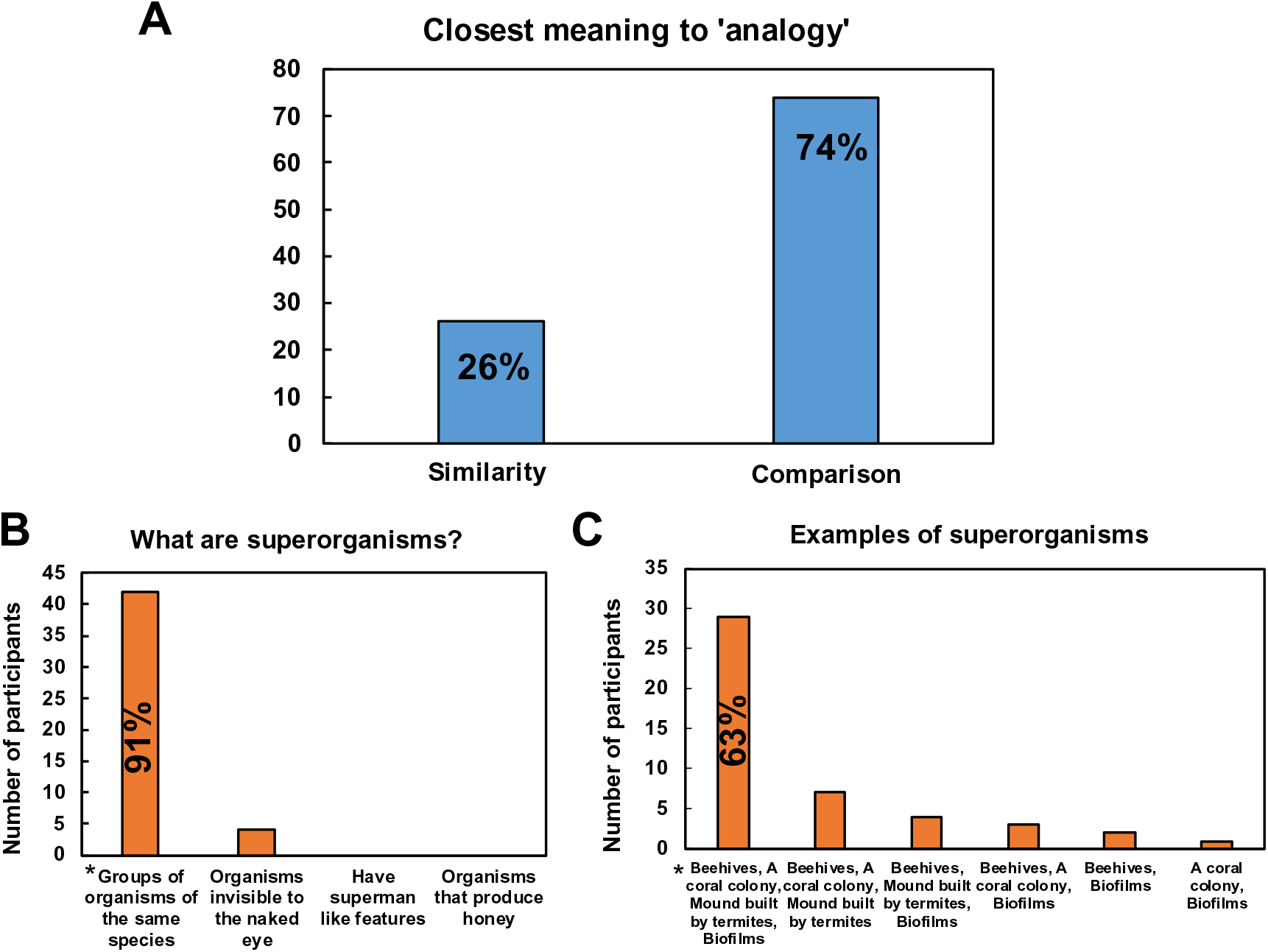
Post-session feedback of the participants on important concepts such as analogies and superorganisms. Feedback collected after the session demonstrates a strong understanding of the basic concept of analogies and superorganisms, which are central to the lesson (* indicates the correct response to the question) (*n=46)*.

**Table 2:** Select anecdotal feedback in response to the question ‘List one way in which beehives and biofilms are similar’.

| **Respondent** | **Feedback** |
| --- | --- |
| **13 years, 8<sup>th</sup> grade** | “They both house multiple members of a species” |
| **13 years, 8<sup>th</sup> grade** | “They both are collections of certain organisms” |
| **14 years, 8<sup>th</sup> grade** | “They both are groups of some living thing that have some special characteristics in common if they are in a group” |
| **20 years, undergraduate** | “Biofilms and beehives are both collections of individual organisms that live together in a structured manner with division of labour and emergent properties” |
| **20 years, undergraduate** | “Both form 3D structure and show division of labor within colonies” |

**Table 5:** Select anecdotal feedback related to the engaging and informative aspects of the analogy-based session.

| **Respondent** | **Feedback** |
| --- | --- |
| **13 years, 7<sup>th</sup> grade** | “The interactive session, the way we imagined different things and how we learnt in a fun way” |
| **15 years, 9<sup>th</sup> grade** | “We ended up learning about two things at the same time. It's easy to correlate between the two topics because we have better understanding of one of the two topics. We could think about a wider range of aspects and come up with better questions” |
| **20 years, undergraduate** | “It used a phenomenon that we are familiar with and could therefore easily relate to. The information was engaging and well placed to ensure that our attention didn't waver too much. The method of explaining was very simple and to the point” |
| **20 years, undergraduate** | “It was more engaging as I was trying to find similarities and difference between two subjects that were compared. It made the understanding of biofilms much easier and interesting” |

### Safety Issues

There are no safety issues associated with the delivery and adoption of this lesson.

## DISCUSSION

### Field Testing

The analogy-based instructional tool was delivered on two separate occasions (session time 90 minutes) via a virtual format (webinar) to high school students (24 students; 13-18 years old) and undergraduates (25 students) across India. While the content and delivery were prepared with these groups in mind, the session for school children was open to younger age groups. High school students were from different schools across the country, and represented a range of grades. The undergraduates were in year 3 of an integrated Masters’ course in Biotechnology, and had completed basic courses in biology and microbiology prior to this session. Registered participants were provided with instructions and pre-session reading materials via email one week before the session. Explicit written participant consent, or parental consent (in the case on participants under 18 years of age), was obtained prior to the collection of feedback.

### Evidence of student learning

Post-session feedback was obtained from 46 respondents, as compared with 49 respondents in the pre-session feedback. It is unclear as to why 3 respondents did not respond or may have left the session prior to completion.

Based on feedback after the session, 74% (n=34/46) students reported the closest meaning of the term ‘analogy’ as comparison and 26% as similarity (Figure 8A). This is in contrast to feedback obtained prior to the session, and indicates a better understanding of analogies as comparative tools. With respect to a general understanding of superorganisms, 91% (n=42/46) of participants identified the correct definition of superorganisms, and 63% (n=29/46) of participants were able to identify all the four examples of superorganisms provided (Figure 8B and C).

Based on feedback, 100% of participants correctly identified biofilms as bacterial communities (Figure 9A), and 70% (n=32/46) answered the importance of studying biofilms as to understand their roles in infection and the environment, and to fight antibiotic resistance (Figure 9B). However, 13% (n=6/46) of participants included the additional option of ‘to learn about beehives’ in their response (Figure 9B). While this was not the response we expected, it does serve to indicate that the concept of using analogies to foster bidirectional understanding of the entities under discussion. In response to the question related to differentiating biofilms from single-celled bacterial forms, 37% (n=17/46) of the participants identified the features of biofilms, namely increased tolerance to antibiotics, groups of microbes and difficulty in biofilm removal, correctly (Figure 9C). On the other hand, only 26% (n=12/46) of participants were also able to identify that the single-celled form of microbial life is less commonly observed as compared with biofilms (Figure 9C).

**Figure 9:**
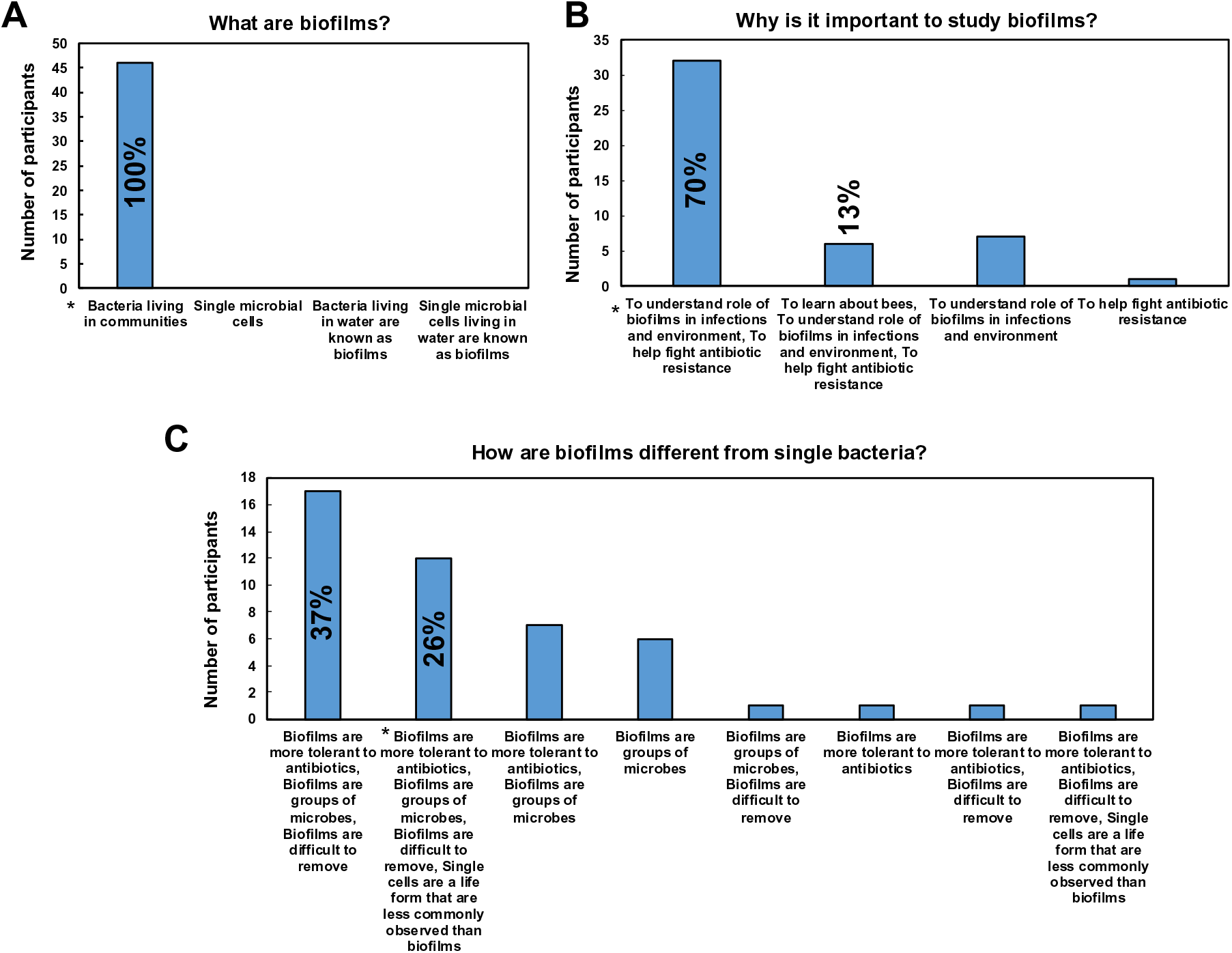
Post-session feedback of the participants on the overall concept of biofilms. Feedback collected after the session demonstrates a strong understanding of the overarching concept of biofilms lesson (* indicates the correct response to the question) (*n=46)*.

Based on the segment on development and structure of biofilms, 80% (n=37/46) of respondents identified ‘attachment’ as the typical first step in biofilm formation (Figure 10A). A total 98% (n=45/46) of participants identified the role of surface properties and cell-to-cell adhesion as important for biofilm formation; with 59% (n=27/46) identifying both (Figure 10B). Further, 96% of participants correctly selected the description of EPS in biofilm matrix (n=44/46). In the segment on chemical communication, 76% (n=35/46) and 72% (n=33/46) of participants correctly identified quorum sensing as the term for bacterial cell density dependent communication and autoinducers as chemical mediators respectively (Figure 10C and D). It is important to note that in the question on chemical communication, ‘pheromones’ was the second most common response (24%, n=11/46), underscoring the importance of highlighting that pheromones are beehive communication molecules. The segment on division of labor in biofilms was largely focused on examples from *B. subtilis* biofilms, with examples drawn from contemporary scientific literature. Based on feedback, 87% (n=40/46) of participants identified the two major matrix components resulting from division of labor in *B. subtilis* biofilms (Figure 10F). However, only 43% (n=20/46) were able to parse out the fact that absence of either one of these matrix components was observed to result in absence of biofilm formation, and 22% (n=10/46) and 35% (n=16/46) answered incorrectly that lack of one component would lead to thicker or normal biofilms respectively (Figure 10G). This is possibly due to the more complex nature of concepts in this segment of the analogy, and subsequent deliveries could focus on clarifying these aspects. In the final segment based on emergent properties, 83% (n=38/46) of participants correctly identified them as properties arising from groups of populations (Figure 10H). This is important to note that in the pre-session feedback, only 56% (n=27/49) had reported being familiar with the term superorganisms (Figure 6A).

**Figure 10:**
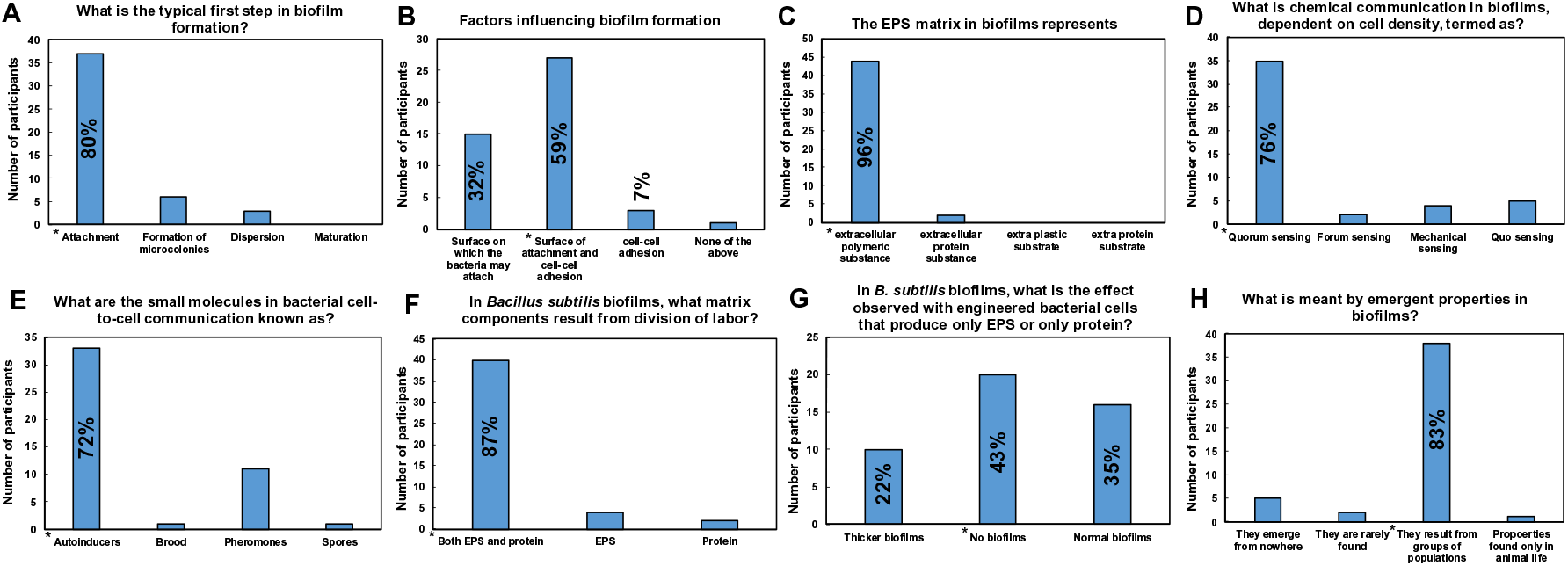
Post-session feedback of the participants on the specific biofilm features covered in this analogy-based lesson. Feedback collected after the session demonstrates an understanding of key features of biofilms, which is in line with the specific learning objectives of the analogy-based lesson (* indicates the correct response to the question) (*n=46)*.

Based on anecdotal feedback, several participants were successfully able to recapitulate similarities between biofilms and beehives (Table 2). Importantly, based on responses, participants were able to highlight differences between biofilms and beehives (Table 3). Several important differences stated by the participants (Table 3) indicate that the differences between the two entities was appreciated. Anecdotal feedback on new ideas that could be explored indicate that the participants were able to leverage comparisons between the two entities to develop novel ideas and lines of investigation for biofilms, not explicitly described in the session (Table 4).

**Table 3:** Select anecdotal feedback in response to the question ‘List one way in which beehives and biofilms are different’.

| **Respondent** | **Feedback** |
| --- | --- |
| **15 years, 9<sup>th</sup> grade** | “Biofilms can be polymicrobial while beehives consist of bees of the same species” |
| **19 years, undergraduate** | “The foraging bees leave and return to the beehive, for collection of nectar and pollen, hence food is not synthesised in the beehive. Whereas, in biofilms, the dispersed bacteria never return to the biofilm and nutrition is synthesised within the colony” |
| **19 years, undergraduate** | “Mode of reproduction” |
| **20 years, undergraduate** | “Beehive has a central figure as the queen bee. Biofilm has no central coordinating element” |

**Table 4:**
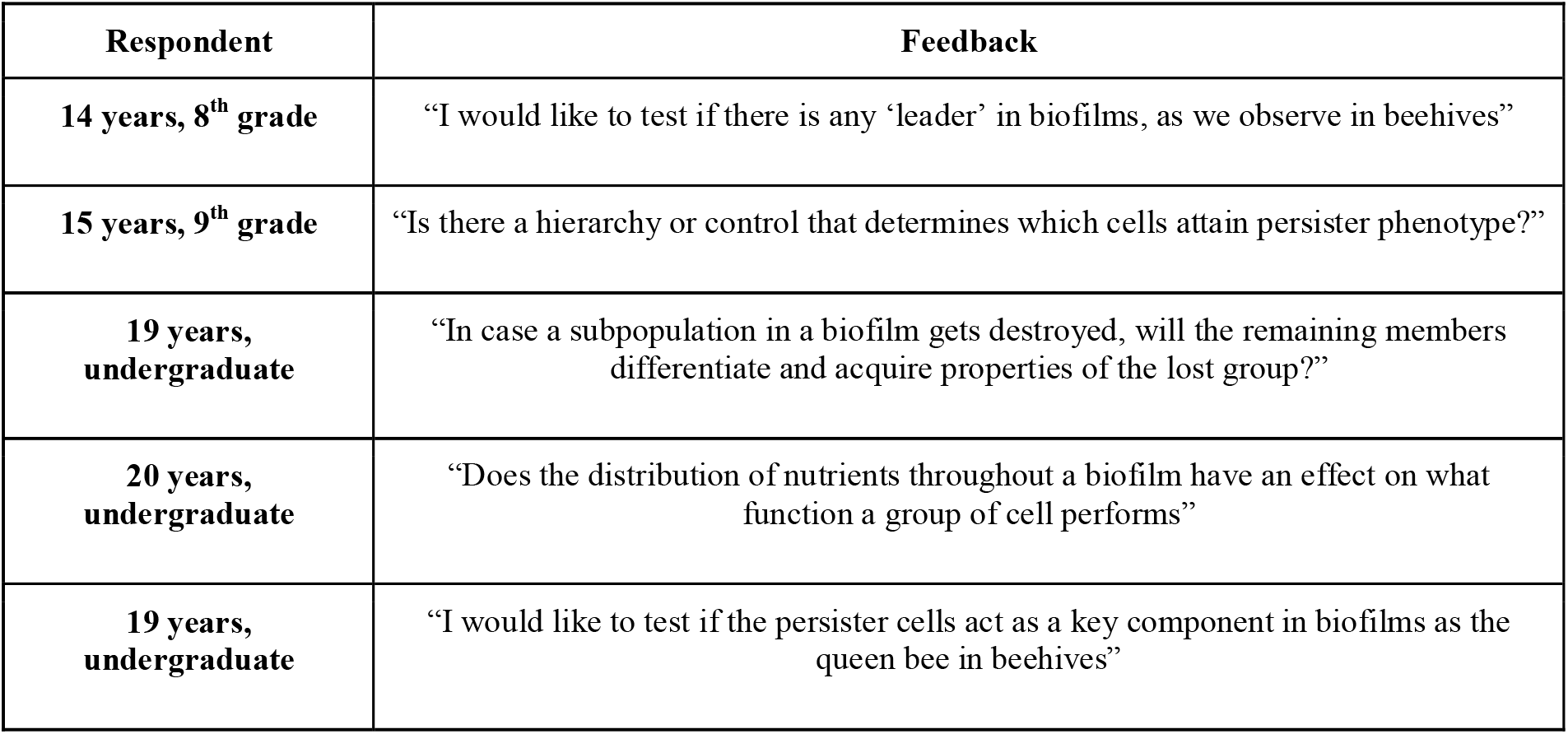
Select anecdotal feedback in response to the question ‘Based on this analogy what new ideas could be explored in biofilms?’, indicating that the analogy can lead to the development of new ideas and hypotheses.

Post-session feedback on the fun, engaging and informative components of the analogy-based instructional session, as well as the level of content in the analogy is shown in Figure 11 and Table 5. When this data was analysed across the two groups, high school students and undergraduates, we did not observe a difference in the responses to the level of content. While this indicates, it was appropriate for both educational groups, depending on the educational level and prior knowledge of the participants, the content of the analogy can be scaled up for undergraduates based on suggestions provided (Suppl Material). Further, 100% of participants responded that they would recommend the analogy-based learning tool to students and teachers for implementation in the curriculum.

**Figure 11:**
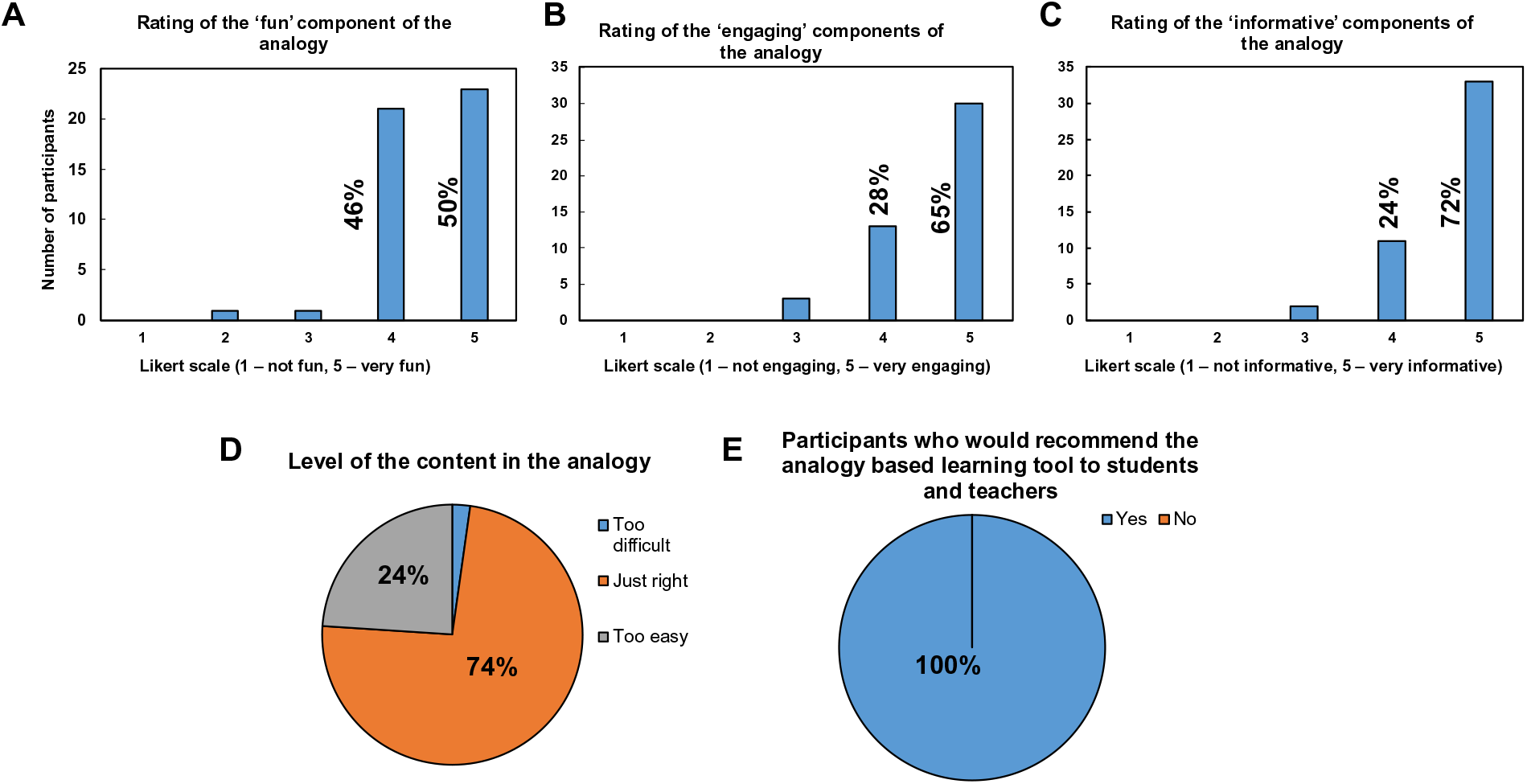
Post-session feedback of the participants regarding the overall learning experience of the session. Based on feedback, the session was reported to be fun, engaging and informative. While majority of the participants reported the level of the content as ‘just right’, based on educational level and prior knowledge, the content can be scaled up with additional suggestions provided (*n=46)*.

### Possible modifications

Based on virtual (webinar-based) delivery and feedback, this analogy-based instructional tool is an effective and engaging approach to introduce the concept of biofilms to high school and undergraduate students. The instructional tool can easily be adapted to in-person delivery with additional group activities. For school classes, this could include group enactment of biofilm formation with assigned roles in the form of name badges or placards. For undergraduates, the lesson could be modified to enable hands on activities where students could use internet resources or a chemistry textbook to illustrate chemical structures of the relevant molecules, with special emphasis on the functional groups. Undergraduates could also be given ‘critical thinking questions’ to work on in groups, such as the advantages of division of labor and emergence in biofilms. To pace the instructional lesson, particularly for classes with students of various learning levels, the instructional tool could also be delivered over two sessions of 1 hour each with discussion time included between the four module segments.

## Supporting information

Slides for delivery

Post-session feedback

Pre-session feedback

Guidelines for Delivery

Key Words

## Acknowledgements

KSK is supported by the Ramalingaswami Re-entry Fellowship (BT/HRD/16/2005), Department of Biotechnology, Government of India. SK was supported by the Ramalingaswami Re-entry Fellowship for part of the duration of this work. We have no conflicts of interest to declare.

## Data availability statement

Data generated in this study, including anonymized feedback data, will be made available by the authors’ upon request.

## Author Contributions

SK, AC, KSK developed the analogy and wrote the manuscript. AC and KSK prepared the analogy-related figures. SK and KSK delivered the analogy-based instructional tool, analysed the feedback data, and prepared the feedback data related figures.

## Notes

**Source of support:** KSK is supported by the Ramalingaswami Re-entry Fellowship (BT/HRD/16/2005), Department of Biotechnology, Government of India. SK was supported by the Ramalingaswami Re-entry Fellowship for part of the duration of this work.

### Competing Interest Statement

The authors have declared no competing interest.

## References

1. Donlan RM, Costerton JW. 2002. Biofilms: Survival mechanisms of clinically relevant microorganisms. Clin Microbiol Rev.

2. Flemming HC, Wingender J, Szewzyk U, Steinberg P, Rice SA, Kjelleberg S. 2016. Biofilms: An emergent form of bacterial life. Nat Rev Microbiol.

3. Madsen JS, Sørensen SJ, Burmølle M. 2018. Bacterial social interactions and the emergence of community-intrinsic properties. Curr Opin Microbiol.

4. Flemming HC, Neu TR, Wozniak DJ. 2007. The EPS matrix: The “House of Biofilm Cells.” J Bacteriol.

5. Rao V, Ghei R, Chambers Y. 2005. Biofilms Research—Implications to Biosafety and Public Health. Appl Biosaf 10:83–90.

6. James GA, Swogger E, Wolcott R, Pulcini E deLancey, Secor P, Sestrich J, Costerton JW, Stewart PS. 2008. Biofilms in chronic wounds. Wound Repair Regen 16:37–44.

7. Motta JP, Wallace JL, Buret AG, Deraison C, Vergnolle N. 2021. Gastrointestinal biofilms in health and disease. Nat Rev Gastroenterol Hepatol 18:314–334.

8. Das S, Singh S, Matchado MS, Srivastava A, Bajpai A. 2019. Biofilms in human health, p. 27–42. In Biofilms in Human Diseases: Treatment and Control. Springer International Publishing.

9. Srivastava S, Bhargava A. 2016. Biofilms and human health. Biotechnol Lett. Springer Netherlands.

10. Murthy PS, Venkatesan R. 2008. Industrial Biofilms and their Control, p. 65–101. In Marine and Industrial Biofouling. Springer Berlin Heidelberg.

11. Di Pippo F, Di Gregorio L, Congestri R, Tandoi V, Rossetti S. 2018. Biofilm growth and control in cooling water industrial systems. FEMS Microbiol Ecol. Oxford University Press.

12. Redelman C V., Marrs K, Anderson GG. 2012. Discovering Biofilms: Inquiry-Based Activities for the Classroom. Am Biol Teach 74:305–309.

13. Wilson EO. 2014. Life on Earth (Unit 1). Apple Books. Wilson Digital, Inc.

14. Fowler S, Rouch R, Wise J. 2013. Concepts of BiologyOpenStax. OpenStax.

15. Weller CM. 1970. The role of analogy in teaching science. J Res Sci Teach 7:113–119.

16. Treagust DF, Harrison AG, Venville GJ. 1998. Teaching Science Effectively With Analogies: An Approach for Preservice and Inservice Teacher Education. J Sci Teacher Educ. Springer.

17. Heywood D. 2002. The place of analogies in science education. Cambridge J Educ 32:233–247.

18. Coll RK, France B, Taylor I. 2005. The role of models/and analogies in science education: Implications from research. Int J Sci Educ. Taylor and Francis Ltd .

19. Brown S, Salter S. 2010. Analogies in science and science teaching. Am J Physiol - Adv Physiol Educ https://doi.org/10.1152/advan.00022.2010.

20. Treagust DF, Duit R, Joslin P, Lindauer I. 1992. Science teachers’ use of analogies: Observations from classroom practice. Int J Sci Educ 14:413–422.

21. Harrison AG, Treagust DF. 1993. Teaching with analogies: A case study in grade-10 optics. J Res Sci Teach 30:1291–1307.

22. Thiele RB, Venville GJ, Treagust DF. 1995. A comparative analysis of analogies in secondary biology and chemistry textbooks used in Australian schools. Res Sci Educ 25:221–230.

23. Jarman R. 1996. Student teachers’ use of analogies in science instruction. Int J Sci Educ 18:869–880.

24. Kipnis N. 2005. Scientific analogies and their use in teaching science, p. 199–233. In Science and Education. Springer.

25. Raimi KT, Stern PC, Maki A. 2017. The Promise and Limitations of Using Analogies to Improve Decision-Relevant Understanding of Climate Change. PLoS One 12:e0171130.

26. Harrison AG, Treagust DF. 2006. Teaching and Learning with Analogies, p. 11–24. In Metaphor and Analogy in Science Education. Springer-Verlag.

27. Glynn SM. 1994. Teaching Science With Analogies A Strategy for Teachers and Textbook Authors.

28. Maharaj-Sharma R, Sharma A. 2015. Observations from secondary school classrooms in Trinidad and Tobago☐: Sci ence teachers ’ use of analogies. Sci Educ Int.

29. Moritz RFA, Southwick EE. 1992. Bees as Superorganisms☐ an Evolutionary Reality. Springer Berlin Heidelberg.

30. Lévesque CM. 2018. Bacterial biofilm☐: the new superorganism. Res Featur.

31. Nowogrodzki R. 1984. Division of labour in the honeybee colony: A review. Bee World.

32. Robinson GE. 1992. Regulation of Division of Labor in Insect Societies. Annu Rev Entomol 37:637–665.

33. Seeley TD. 1995. The Wisdom of the Hive: The Social Physiology of Honey Bee Colonies. Harvard University Press.

34. Nazzi F. 2016. The hexagonal shape of the honeycomb cells depends on the construction behavior of bees. Sci Rep 6.

35. Seeley T. 1977. Measurement of nest cavity volume by the honey bee (Apis mellifera). Behav Ecol Sociobiol https://doi.org/10.1007/BF00361902.

36. Seeley TD, Morse RA. 1978. Nest site selection by the honey bee, Apis mellifera. Insectes Soc https://doi.org/10.1007/BF02224297.

37. Schmidt JO, Thoenes SC. 1992. Criteria for nest site selection in honey bees (Hymenoptera: Apidae)☐: preferences between pheromone attractants and cavity shapes. Environ Entomol https://doi.org/10.1093/ee/21.5.1130.

38. Barron AB, Plath JA. 2017. The evolution of honey bee dance communication: A mechanistic perspective. J Exp Biol.

39. Seeley TD, Visscher PK. 2003. Choosing a home: How the scouts in a honey bee swarm perceive the completion of their group decision making. Behav Ecol Sociobiol https://doi.org/10.1007/s00265-003-0664-6.

40. Seeley TD, Buhrman SC. 1999. Group decision making in swarms of honey bees. Behav Ecol Sociobiol https://doi.org/10.1007/s002650050536.

41. Stoodley P, Sauer K, Davies DG, Costerton JW. 2002. Biofilms as complex differentiated communities. Annu Rev Microbiol 56:187–209.

42. Sifri CD. 2008. Quorum sensing: Bacteria talk sense. Clin Infect Dis https://doi.org/10.1086/592072.

43. Limoli DH, Jones CJ, Wozniak DJ. 2015. Bacterial Extracellular Polysaccharides in Biofilm Formation and Function. Microbiol Spectr https://doi.org/10.1128/microbiolspec.mb-0011-2014.

44. Hall-Stoodley L, Stoodley P. 2002. Developmental regulation of microbial biofilms. Curr Opin Biotechnol 13:228–233.

45. Bridier A, Dubois-Brissonnet F, Boubetra A, Thomas V, Briandet R. 2010. The biofilm architecture of sixty opportunistic pathogens deciphered using a high throughput CLSM method. J Microbiol Methods 82:64–70.

46. Shrout JD, Tolker-Nielsen T, Givskov M, Parsek MR. 2011. The contribution of cell-cell signaling and motility to bacterial biofilm formation. MRS Bull 36:367–373.

47. Schuster JJ, Markx GH. 2014. Biofilm architecture. Adv Biochem Eng Biotechnol 146:77–96.

48. Bortolotti L, Costa C. 2014. Chemical communication in the honey bee society, p. 147–210. In Neurobiology of Chemical Communication. CRC Press.

49. AL-Kahtani SN, Bienefeld K. 2012. The Nasonov Gland Pheromone is Involved in Recruiting Honeybee Workers for Individual Larvae to be Reared as Queens. J Insect Behav 25:392–400.

50. Miller MB, Bassler BL. 2001. Quorum sensing in bacteria. Annu Rev Microbiol 55:165–199.

51. Rutherford ST, Bassler BL. 2012. Bacterial quorum sensing: Its role in virulence and possibilities for its control. Cold Spring Harb Perspect Med.

52. Abisado RG, Benomar S, Klaus JR, Dandekar AA, Chandler JR. 2018. Bacterial quorum sensing and microbial community interactions. MBio.

53. Li YH, Tian X. 2012. Quorum sensing and bacterial social interactions in biofilms. Sensors.

54. Hense BA, Schuster M. 2015. Core Principles of Bacterial Autoinducer Systems. Microbiol Mol Biol Rev https://doi.org/10.1128/mmbr.00024-14.

55. Nadell CD, Xavier JB, Levin SA, Foster KR. 2008. The evolution of quorum sensing in bacterial biofilms. PLoS Biol https://doi.org/10.1371/journal.pbio.0060014.

56. Pena RT, Blasco L, Ambroa A, González-Pedrajo B, Fernández-García L, López M, Bleriot I, Bou G, García-Contreras R, Wood TK, Tomás M. 2019. Relationship between quorum sensing and secretion systems. Front Microbiol.

57. Ng W-L, Bassler BL. 2009. Bacterial Quorum-Sensing Network Architectures. Annu Rev Genet https://doi.org/10.1146/annurev-genet-102108-134304.

58. Verbeke F, De Craemer S, Debunne N, Janssens Y, Wynendaele E, Van de Wiele C, De Spiegeleer B. 2017. Peptides as quorum sensing molecules: Measurement techniques and obtained levels in vitro and in vivo. Front Neurosci.

59. Mackensen O. 1951. Viability and Sex Determination in the Honey Bee (Apis Mellifera L.). Genetics 36:500–509.

60. Charlesworth B. 2003. Sex determination in the honeybee. Cell.

61. Gempe T, Hasselmann M, Schiøtt M, Hause G, Otte M, Beye M. 2009. Sex determination in honeybees: Two separate mechanisms induce and maintain the female pathway. PLoS Biol https://doi.org/10.1371/journal.pbio.1000222.

62. Page RE, Robinson GE. 1991. The Genetics of Division of Labour in Honey Bee Colonies. Adv In Insect Phys https://doi.org/10.1016/S0065-2806(08)60093-4.

63. Stewart PS, Franklin MJ. 2008. Physiological heterogeneity in biofilms. Nat Rev Microbiol 6:199–210.

64. van Gestel J, Vlamakis H, Kolter R. 2015. Division of Labor in Biofilms: the Ecology of Cell Differentiation. Microbiol Spectr https://doi.org/10.1128/microbiolspec.mb-0002-2014.

65. Dragos A, Kiesewalter H, Martin M, Hsu C-Y, Hartmann R, Wechsler T, Eriksen C, Brix S, Drescher K, Stanley-Wall N, Kummerli R, Kovacs AT. 2018. Division of Labor during Biofilm Matrix Production. Curr Biol 28:1903–1913.e5.

66. Momeni B. 2018. Division of Labor: How Microbes Split Their Responsibility. Curr Biol.

67. Armbruster CR, Lee CK, Parker-Gilham J, De Anda J, Xia A, Zhao K, Murakami K, Tseng BS, Hoffman LR, Jin F, Harwood CS, Wong GCL, Parsek MR. 2019. Heterogeneity in surface sensing suggests a division of labor in pseudomonas aeruginosa populations. Elife https://doi.org/10.7554/eLife.45084.

68. Ponge J-F. 2005. Emergent properties from organisms to ecosystems: towards a realistic approach. Biol Rev Camb Philos Soc 80:403–411.

69. Delaplane KS. 2017. Emergent Properties in the Honey Bee Superorganism. Bee World https://doi.org/10.1080/0005772x.2017.1290893.

70. Hall CW, Mah T-F. 2017. Molecular mechanisms of biofilm-based antibiotic resistance and tolerance in pathogenic bacteria. FEMS Microbiol Rev 41:276–301.

71. Brown MRW, Allison DG, Gilbert P. 1988. Resistance of bacterial biofilms to antibiotics a growth-rate related effect? J Antimicrob Chemother https://doi.org/10.1093/jac/22.6.777.

72. Grant SS, Hung DT. 2013. Persistent bacterial infections, antibiotic tolerance, and the oxidative stress response. Virulence.

73. Conlon BP, Rowe SE, Lewis K. 2015. Persister cells in biofilm associated infections. Adv Exp Med Biol https://doi.org/10.1007/978-3-319-09782-4_1.

