## Supplementary material for "Of Biofilms and Beehives: An Analogy-Based Instructional Tool to Introduce Biofilms to High-School and Undergraduate Students": Slides for delivery: Slides for delivery.pdf

### What is an 'Analogy'?

A comparison between one thing and another, for the purpose of understanding. Explain one thing in terms of another to highlight the ways in which they are alike.

Can you think of an analogy in everyday life?

### General introduction to beehives

What are 'superorganisms'?

Can you name some?

Can you describe beehives you have seen around you?

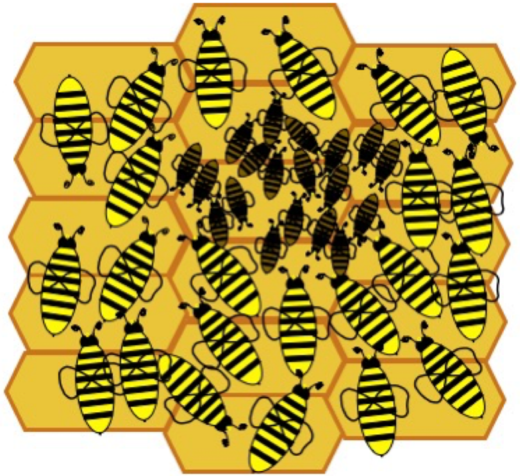

### General introduction to biofilms

What are microbes?

What are biofilms?

Where are biofilms seen in the human body?

Where are biofilms found in the environment?

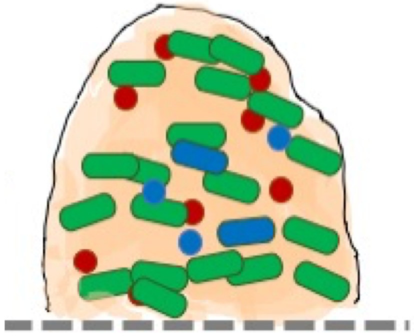

What are the effects of biofilms - harmful and beneficial?

### Using beehives as an analogy to learn about biofilms

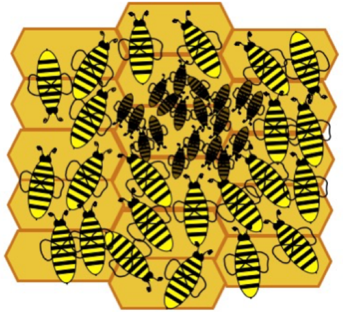

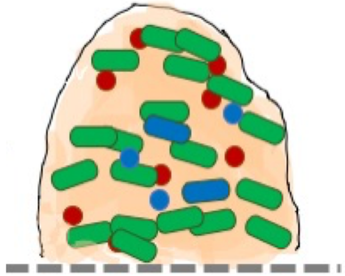

Both are 'superorganisms'

Beehives are visible to the eye, well-studied and commonly observed

Biofilms are microscopic, studied in research laboratories, and have far reaching consequences on human health and the environment

### Using beehives as an analogy to learn about biofilms

Our analogy is based on four segments -

Development and structure of beehives and biofilms
Chemical communication in beehives and biofilms
Division of labor in beehives and biofilms
Emergent properties of beehives and biofilms

### Development and structure of beehives and biofilms

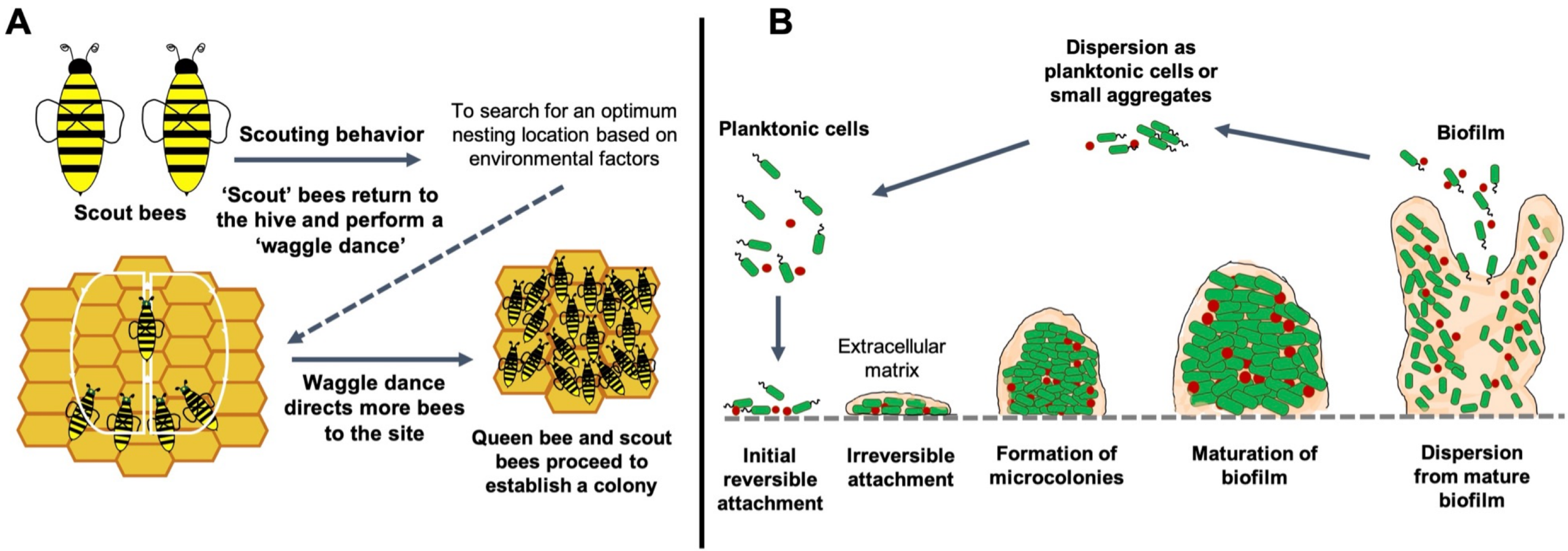

### Guidelines for discussion

We will state the word and you tell us the function/description!

Scouting behavior, waggle dance

EPS, attachment, maturation, 3D structure

### Chemical communication in beehives and biofilms

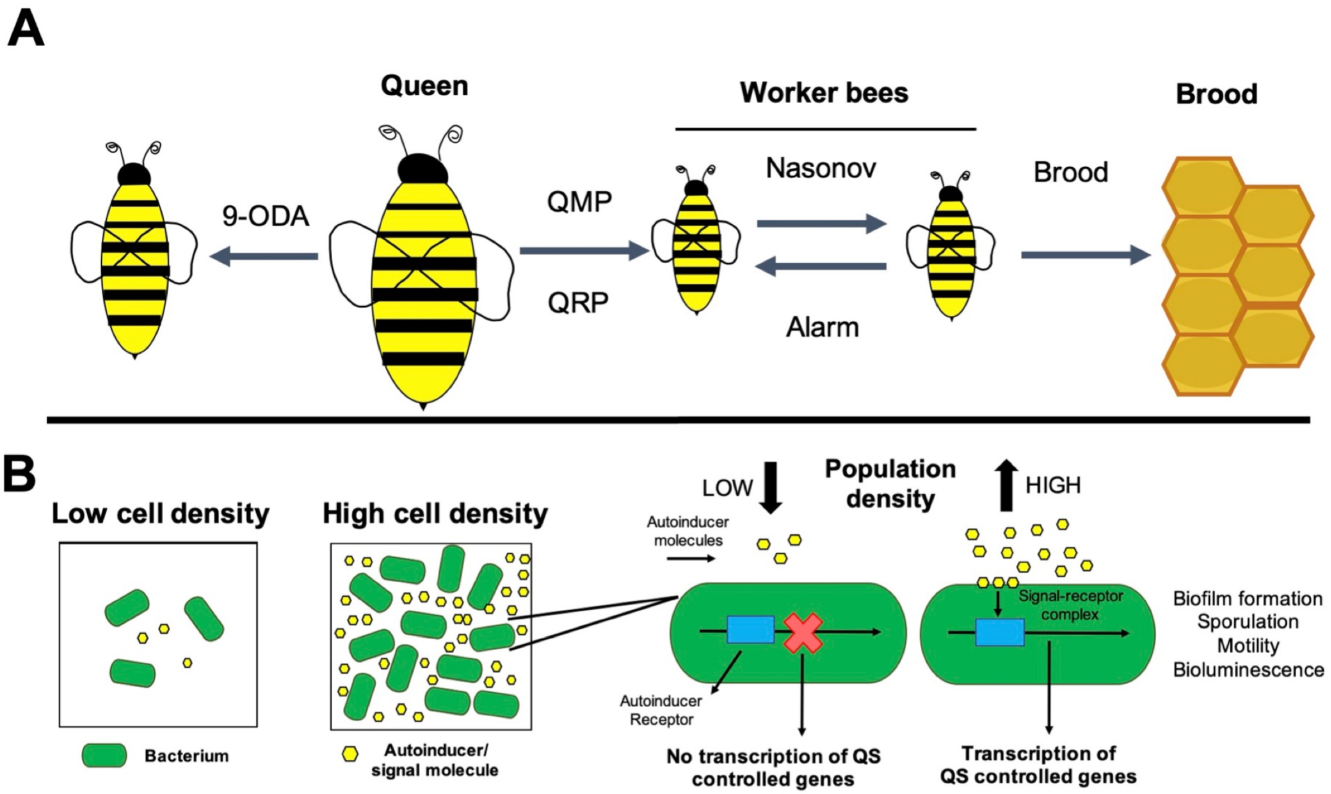

### Guidelines for discussion

What is the role of pheromones in beehive communication?

What is the role of autoinducers in biofilm communication?

Let's name some group behaviours in bacteria!

### Division of labor in beehives and biofilms

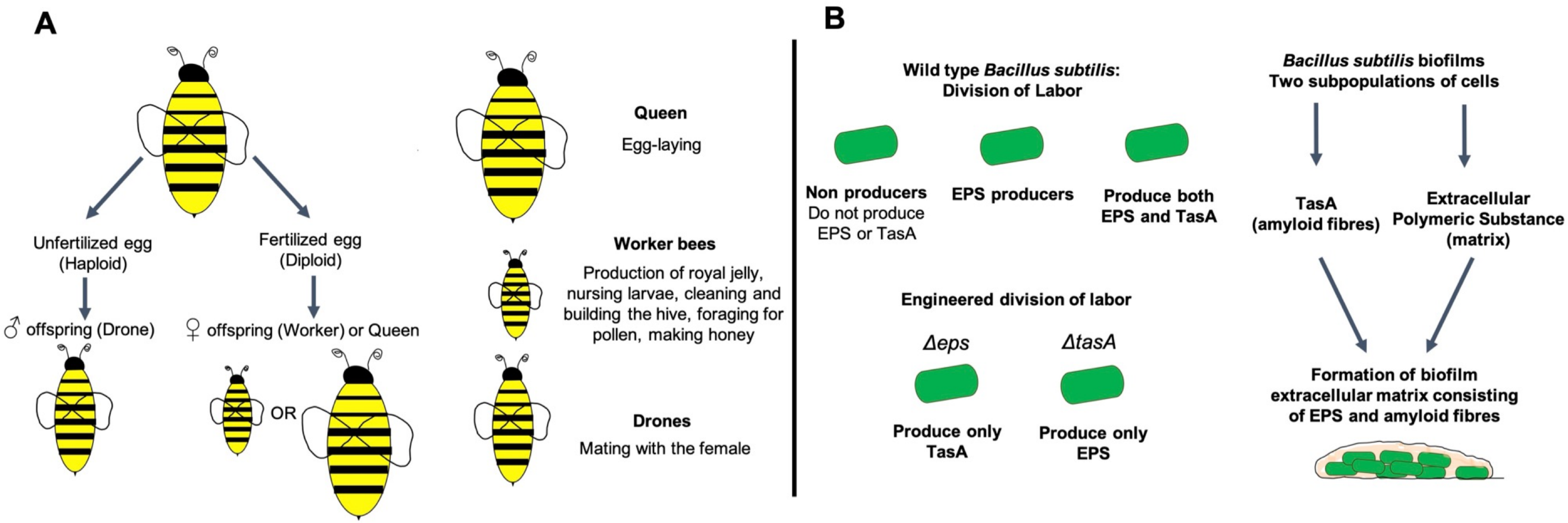

### Guidelines for discussion

Let's play some roles!

We will call out a bee type and you can tell us its functions

We will call out a shared biofilm product and you can tell us its functions

What might be the possible advantages and disadvantages of division of labor in biofilms?

### Emergent properties of beehives and biofilms

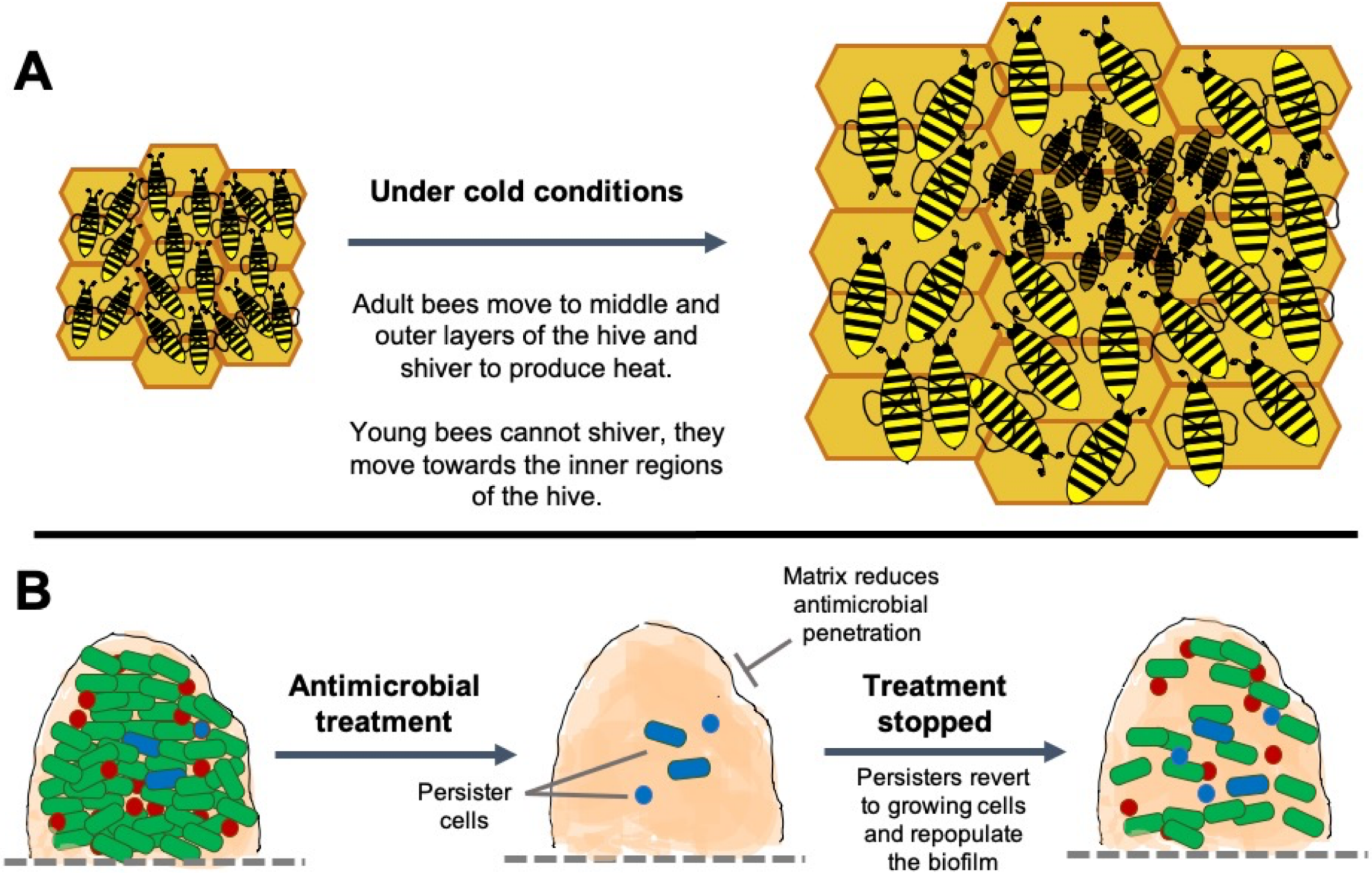

### Guidelines for discussion

Let's think about -

Examples of emergent behavior in nature

Examples of emergent behavior in other animals or insects

Why is thermoregulation beneficial to beehives?

Why is antibiotic tolerance an emergent property of biofilms, and not single cells?

### Limitations and misconceptions

What are some features not shared between beehives and biofilms?

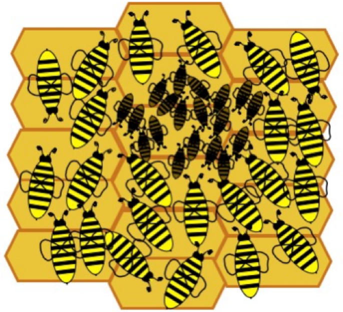

Think -

Organism

Species

structure

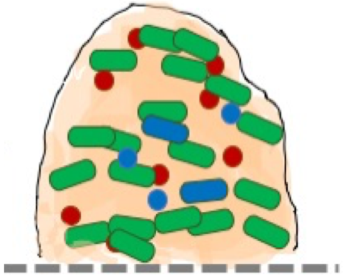

### New ideas and hypotheses

Can you think of new ideas to explore in biofilms based on what we know of beehives?

Think of the 'core element' or the queen in beehives - can you build an idea based on this?

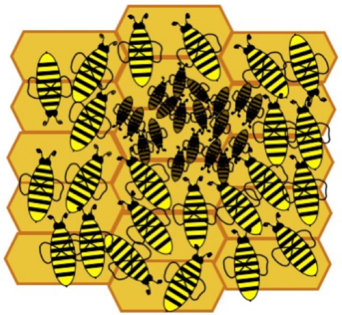

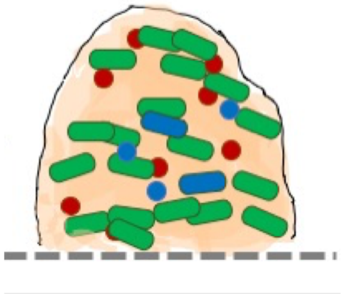

**Pre-session feedback form (filled via Google forms)**

1. **Participant details (to be filled in)**

   Name (optional):

   Age (in years):

   Email ID (optional):

   Gender (female/male/non-binary/prefer not to state):

   Course being pursued (only for undergraduates):

   Education level being pursued (grade or year of education):

2. **Pre-session reading materials (tick the option)**

   Did you use or read the suggested pre-session reading materials? (Yes/No)

   If you did, which materials did you use? (you can tick more than one)

   (Key words provided/basic biology textbooks/the description of the analogy)

3. **General questions about analogies (tick the option)**

   Have you heard of the word 'analogy'? (Yes/No)

   What do you think the closest meaning of the word 'analogy' is?
   (Similarity/Comparison/Contrast/Differences)

   Have you come across analogy based materials or have analogies been used in your curriculum for teaching or educational purposes? (Yes/No)

4. **General questions about beehives and biofilms (tick the option)**

   Have you seen or heard of beehives? (Yes/No)

   Have you heard of biofilms? (Yes/No)

   Have you heard of superorganisms? (Yes/No)

   Do your biology or science textbooks/classes include descriptions or lessons on beehives? (Yes/No)

   Do your biology or science textbooks/classes include descriptions or lessons on microbial life? (Yes/No)

   Do your biology or science textbooks/classes include descriptions or lessons on biofilms? (Yes/No)

   Do your biology or science textbooks/classes include descriptions or lessons on superorganisms? (Yes/No)

**Post-session feedback form (filled via Google forms)**

1. **How useful did you find the pre-session reading materials?**

   1 2 3 4 5

   (not helpful) (very helpful)

2. **Feedback related to analogies and superorganisms**

   What do you think the closest meaning of the word 'analogy' is?
   (Similarity/Comparison/Contrast/Differences)

   In biology, what are superorganisms? (choose one)

   - Have superhuman-like features
   - Interacting groups of organisms of the same species
   - Organisms invisible to the naked eye
   - Defined as organisms that produce honey

   Tick all that are examples of superorganisms (tick all that apply)

   - Beehives
   - A coral colony
   - Mound built by termites
   - Biofilms

3. **Feedback related to general understanding of biofilms**

   What are biofilms?

   - Single microbial cells
   - Bacteria living in water are known as biofilms
   - Bacteria living in communities
   - Single microbial cells living in water are known as biofilms

   Why is it important to study biofilms? (tick all that apply)

   - To learn about bees
   - To understand the role of biofilms in infections and environment
   - To help fight antibiotic resistance

   How are biofilms different from single microbes or bacteria? (tick all that apply)

   - Biofilms are more tolerant to antibiotics
   - Biofilms are groups of microbes
   - Biofilms are difficult to remove
   - Single cells are a life form that are less commonly observed than biofilms

4. **Feedback related to understanding of specific features of biofilms**

   What is the typical first step of biofilm formation?

- Attachment
- Formation of microcolonies
- Dispersion
- Maturation

What are the factors that influence biofilm formation?

- Surface on which the bacteria may attach
- Cell-to-cell adhesion
- Both of the above
- None of the above

The EPS matrix of biofilms is

- Extracellular polymeric substance
- Extra plastic substrate
- Extracellular protein substrate
- Extra protein substrate

What is chemical communication in biofilms, dependent on cell density, known as?

- Quo sensing
- Mechanical sensing
- Quorum sensing
- Forum sensing

What are the small molecules in bacterial cell to cell communication known as?

- Pheromones
- Spores
- Autoinducers
- Brood

In *Bacillus subtilis* biofilms, what matrix components result from division of labor?

- EPS
- Protein
- Both EPS and protein

In *B. subtilis* biofilms, what is the effect observed with engineered bacterial cells that produce only EPS or only protein?

- Thicker biofilms
- No biofilms
- Normal biofilms

What is meant by 'emergent' properties in biofilms?

- Properties that emerge from nowhere
- Properties that are rarely observed
- They result from groups of microbial populations
- They are found only in microbial life

**5. Free response questions related to understanding of specific features of the analogy**

List one way in which biofilms and beehives are similar.

List one way in which biofilms and beehives are different.

Based on this analogy, what new idea or hypothesis you would like to test in biofilms?

**6. Feedback related to the analogy-based learning experience**

On a scale of 1-5, how 'fun' was the analogy?

1 2 3 4 5

(1-lowest, 5-highest)

On a scale of 1-5, how 'engaging' was the analogy?

1 2 3 4 5

(1-lowest, 5-highest)

On a scale of 1-5, how 'informative' was the analogy?

1 2 3 4 5

(1-lowest, 5-highest)

How would you rate the level of difficulty of the analogy content?

- Too easy
- Just right
- Too difficult

Would you recommend this analogy-based learning tool to students and teachers in your school or university? (Yes/No)

**Guidelines for delivery of the analogy**

1. **Pre-session feedback – 10 minutes** (as in provided feedback form)

2. **General introduction to analogies (5 minutes)**

- Description of the term analogy
- Examples of analogies in everyday life and science

3. **General introduction to beehives (5 minutes)**

- Introduction to 'superorganism states', with examples drawn from student observations and experiences (ant colonies, sponges, schools of fish, beehives)

- Descriptions of beehives observed by students (locations, observations of bee behaviours, hive structure)

4. **General introduction to biofilms (5 minutes)**

- Discussion on microbial communities (distinction from single-celled forms, different microbial species can coexist in communities, play a key role in health and environmental processes)

- Key features of organized microbial communities or 'biofilms' (bacterial cells as aggregates, presence of a matrix or 'glue', coexistence of different species of bacteria)

- Examples of biofilms in the human body (dental plaque, gut bacteria, biofilms that form on prosthetic devices such as catheters, valves, replacement joints), and effects of biofilms on human health (persistent infections, long term antibiotic treatments)

- Examples of biofilms in the environment (ship hulls, pond scum, sewage reactors), and effects of biofilms on industry and environment (pipe blockages, water-logging, reduction of equipment efficiency)

- Biofilms can also serve beneficial functions, whereby bacteria in biofilms or biofilm products such as enzymes, can be used for food and agriculture applications such as food fermentation, biofertilizers, wastewater treatment (Turhan et al., 2019)

**5. Development and structure of beehives and biofilms (10 minutes)**

For this section, the description of the analogy and Figure 1 is to be used to share and discuss key points related to the development and structure of beehives and biofilms.

**Suggested additional activities:** In an in-person format, the instructor could divide the class into two groups (or multiple groups, depending on class size) for an interactive activity based on development and structure of beehives and biofilms. For school classes, this could be accompanied by assigned roles (scout bees, planktonic cells, matrix) with name badges or placards. This could also include group enactment of beehive formation with features such as scouting behaviour, waggle dance, recruitment dance, followed by colony formation or group enactment of biofilm formation, starting with the initial attachment of few single cells, followed by irreversible attachment, cell-to-cell signalling, EPS production, and biofilm maturation to form a heterogeneous structure.

**6. Chemical communication in beehives and biofilms (10 minutes)**

For this section, the description of the analogy and Figure 2 is to be used to share and discuss key points related to chemical communication in beehives and biofilms.

**Suggested additional activities:** Following this section of the analogy, students could work in small groups for a discussion, followed by a hands-on activity on the chemical substances that mediate communication in beehives and biofilms. For this, they could use internet resources or a chemistry textbook to illustrate chemical structures of the relevant molecules, with special emphasis on the functional groups. Alternatively, the role and functioning of the chemical molecules could be discussed, and students could turn in the illustrated chemical structures as an assignment.

Communication in beehives

- Differences between volatile and non-volatile chemicals (structural and functional level differences)

- Role of pheromones in animal communication, with a focus on insect communication

- Illustrate chemical structures of volatile pheromones, non-volatile pheromones, Queen Mandibular Pheromone (QMP), Nasonov gland pheromones, alarm pheromones, drone pheromones

Communication in biofilms

- Discuss the phenomenon of density-dependent bacterial communication, and its effects on changes in group functions (biofilm formation, motility, sporulation)

- Role of autoinducers in bacterial communication, with a focus on intraspecies and interspecies autoinducers

- Illustrate chemical structures of the three major classes of autoinducer molecules, N-acylated homoserine lactones (AI-1), oligopeptides and the interspecies chemical signal, autoinducer-2 (AI-2, furanosyl borate diester)

**7. Division of labor in beehives and biofilms (10 minutes)**

For this section, the description of the analogy and Figure 3 is to be used to share and discuss key points related to division of labor in beehives and biofilms.

In this segment, the instructor and students could work to understand the examples of division of labor stated in the analogy, and develop insights into the advantages and possible disadvantages of labor segregation in collective groups.

- Discuss the roles played by different bee subpopulations, and how this could be advantageous to the beehive as a whole (cooperation, efficiency, survival, resilience)

- Discuss the collective benefits of shared products in biofilm structures (nutrients, enzymes, metabolic factors, matrix proteins)

**Suggested additional activities:** At the end of this discussion students can be given a 'critical thinking question', that they can work in groups to share ideas, and prepare a written response. An example of such a question is, 'What might be the possible disadvantages of division of labor and shared factors in biofilm populations?' Following this, students can be directed to additional reading resources that discuss the implications of division of labor in microbial populations, such as long term trade-offs and emergence of cheater populations.

**8. Emergent properties in beehives and biofilms (10 minutes)**

For this section, the description of the analogy and Figure 4 is to be used to share and discuss key points related to emergent properties of beehives and biofilms.

The discussion could be focused on specific examples of emergent properties in beehives and biofilms.

- How does thermoregulation serve as an example of an emergent property in beehives, and why is this a collective behaviour?

- Why is antibiotic tolerance an emergent property in multicellular microbial communities such as biofilms, and why is this distinctly different from single-celled microbial forms?

**Suggested additional activities:** In the final segment of the analogy, the discussion could be focused on the phenomenon of emergence, and understanding emergent behaviour, and examples from physical and biological systems (snowflakes, ant colonies, schools of fish)

**9. Limitations of this analogy, and possible misconceptions (5 minutes)**

To address this, students can be recommended to highlight features not shared between biofilms and beehives. For example, the most fundamental difference between the two states is that bacteria are prokaryotes whereas bees are eukaryotes. Another important difference lies in the species-level composition of biofilms and beehives. Beehives most often comprise a single-species of honeybees, and owing to species-specific pheromones interspecies hives are typically not encountered in nature (Su et al., 2008). Biofilms, on the other hand, most often consist of multiple bacterial species (or polymicrobial biofilms), and can also comprise bacterial and fungal species (Willems et al., 2016). Along these lines, other specific differences related to development, structure, communication, and emergent properties can be discussed. For example, when students picture the 'wiggle dance', they could have an impression of a monolayer of bees. However, a fully-formed biofilm is not a monolayer, but is in fact a heterogeneous, three-dimensional structure containing aggregates of bacterial cells separated by water and nutrient channels. Further, the emergent functions in biofilms and beehives are different, in that the bees have an 'active' form or response (for example for thermoregulation) whereas in the biofilm, it is mainly 'passive' in that most of the properties result from a static physical barrier (antibiotic tolerance).

**10. Can this analogy lead to new ideas and hypotheses? (5 minutes)**

In this segment, the students can be suggested to explore if the instructional analogy with beehives could enable the development of new ideas related to biofilms. For this, the instructor can present certain case scenarios, or the students could work in groups, to identify and discuss the possibility of certain features of beehives leading to potential lines of biofilm investigation. For example, in beehives there exists a 'core' element (the queen) that serves to control, coordinate and orchestrate the functioning of the hive. Based on this, a question can arise as to whether there is an equivalent 'core' group of bacteria in the biofilm that is critical to its survival? This has implications, as if we were to identify such a core group, this group could be the target for biofilm removal strategies. This could lead to the discussion of the role of specialized cell groups such as dormant cells and persisters, widely considered to contribute to the recalcitrance and recurrence of biofilms (Lewis, 2005).

11. **Post-session feedback – 15 minutes** (as in provided feedback form)

**Key words for suggested reading prior to the analogy lesson on biofilms**

| | | | | |
|---|---|---|---|---|
| Beehives | Superorganisms | Emergent properties | Scouting behaviour | Waggle dance |
| | Chemical communication | Pheromones | Exocrine glands | Volatile and non-volatile chemicals |
| | Odorant receptors | Division of labor | Haplodiploid mode of sex determination | Thermoregulation |
| Biofilms | Microbes | Microbial growth | Van der Waals interactions | Hydrophobic interactions |
| | Cell-to-cell communication | Quorum sensing | Autoinducers | Gene expression |
| | Mutations | Antibiotics and antimicrobials | Antibiotic tolerance | Persisters |

1.
